# Mechanism of DNA entrapment by the MukBEF SMC complex and its inhibition by a viral DNA mimic

**DOI:** 10.1101/2024.10.02.616235

**Authors:** Frank Bürmann, Bryony Clifton, Sophie Koekemoer, Oliver J. Wilkinson, Dari Kimanius, Mark S. Dillingham, Jan Löwe

## Abstract

Ring-like structural maintenance of chromosomes (SMC) complexes are crucial for genome organization and operate through mechanisms of DNA entrapment and loop extrusion. Here, we explore the DNA loading process of the bacterial SMC complex MukBEF. Using electron cryomicroscopy (cryo-EM), we demonstrate that ATP binding opens one of MukBEF’s three potential DNA entry gates, exposing a DNA capture site that positions DNA at the open neck gate. We discover that the gp5.9 protein of bacteriophage T7 blocks this capture site by DNA mimicry, thereby preventing DNA loading and inactivating MukBEF. We propose a comprehensive and unidirectional loading mechanism in which DNA is first captured at the complex’s periphery and then ingested through the DNA entry gate, powered by a single cycle of ATP hydrolysis. These findings illuminate a fundamental aspect of how ubiquitous DNA organizers are primed for genome maintenance and demonstrate how this process can be disrupted by viruses.

**Graphical abstract:** 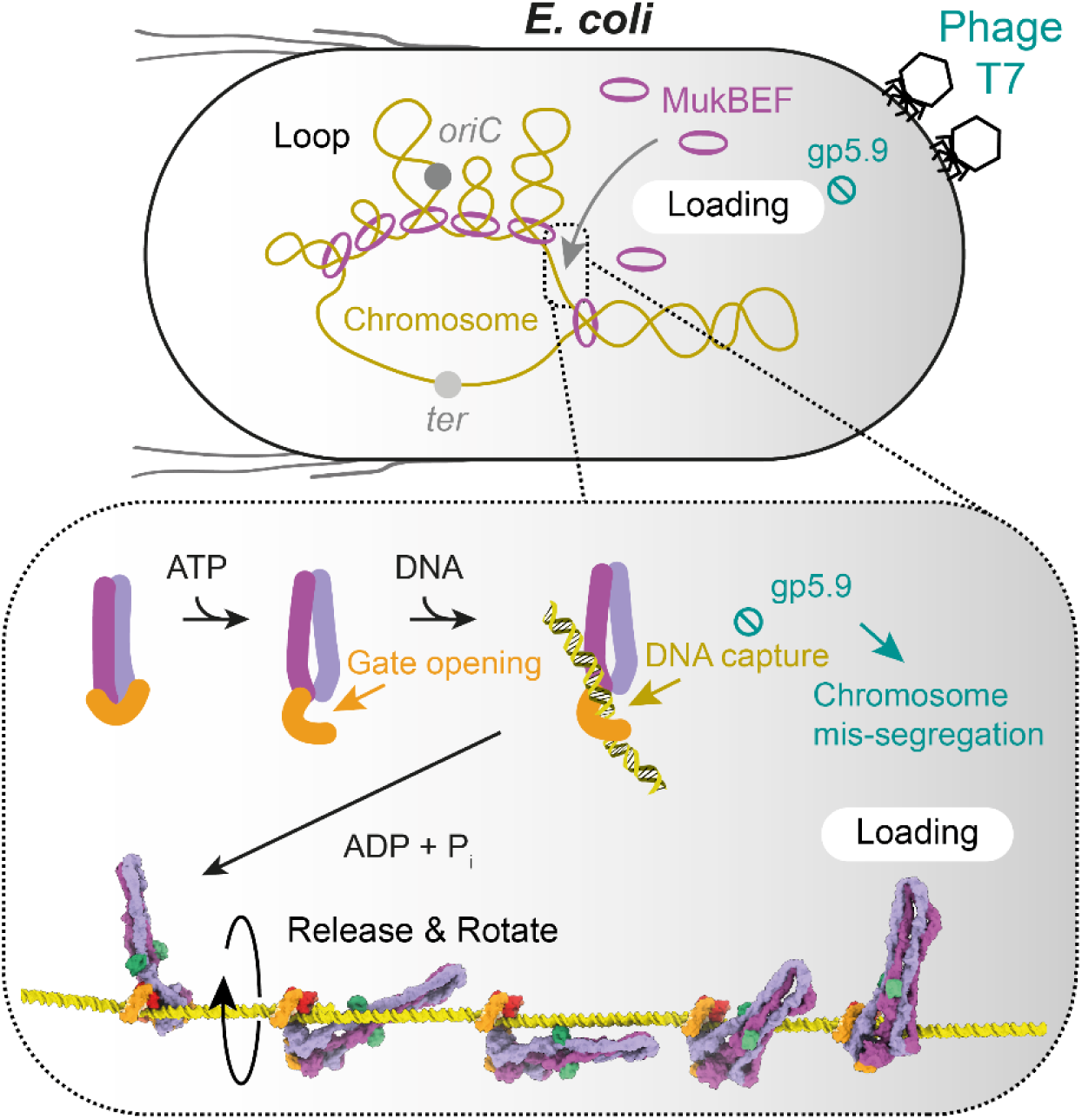

## Introduction

Large ring-like structural maintenance of chromosomes (SMC) complexes are fundamental chromosome organizers, and facilitate diverse DNA transactions in bacteria, archaea, and eukaryotes ^1–3^. They mediate mitotic and meiotic chromosome compaction, sister chromatid cohesion, folding of chromosomes, DNA recombination, double-strand break repair, silencing of viral genomes, and the restriction of plasmids ^4–13^. SMC functions are based on the entrapment of DNA within the complex and the ATP-powered extrusion of large DNA loops. DNA entrapment was first described for its role in sister chromatid cohesion, where replicated sister DNAs are held together by the cohesin complex ^14–16^. However, other SMC complexes entrap DNA without mediating cohesion, suggesting that entrapment has another more fundamental purpose ^17–20^. How exactly entrapment is established by loading DNA into an SMC complex, and how entrapment relates to loop extrusion, is largely unclear.

MukBEF was the first SMC complex discovered, and folds the chromosome of *Escherichia coli* and related bacteria ^8,21,22^. It is a member of the <u>M</u>ukBEF-li<u>k</u>e <u>S</u>MC (Mks) or Wadjet group (**Figure 1A**), many members of which associate with the nuclease MksG/JetD to protect bacteria against plasmid infection ^13,23–26^. MukBEF has a key role in chromosome segregation, and like several other Wadjet group members lacks MksG/JetD ^23^ (**Figure S1A**). MukBEF deficiency is lethal under fast-growing conditions and accompanied by defective chromosome segregation and an increased production of anucleate cells ^21^.

**Figure 1.**
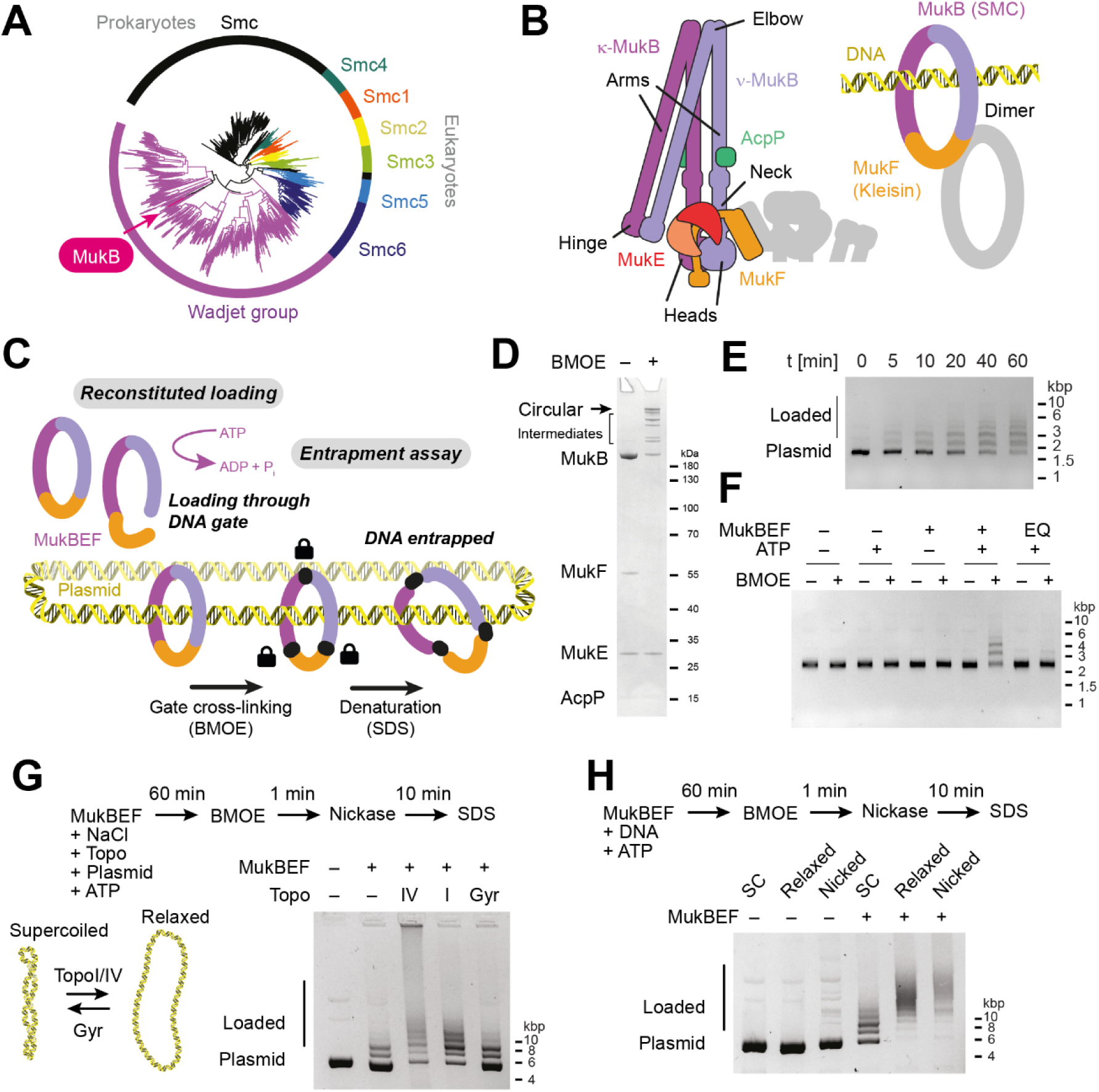
Reconstitution of DNA loading. (**A**) Phylogenetic tree of SMC proteins inferred from chained alignments of head and hinge regions. (**B**) Architecture of MukBEF (left) and simplified geometry of the complexes indicating DNA entrapment (right). (**C**) Concept of the *in vitro* loading assay. MukBEF is loaded onto plasmid DNA in the presence of DNA, then gates are closed by BMOE-mediated cysteine cross-linking, and protein/DNA catenanes are probed after SDS denaturation. (**D**) BMOE cross-linking of *P. thracensis* MukBEF containing cysteine residues in the three gate interfaces. A Coomassie stained SDS-PAGE gel shows cross-linked products. (**E**) Loading time course of MukBEF on negatively supercoiled DNA (pFB527) in the presence of an ATP regeneration system. Reactions were terminated by BMOE cross-linking at the indicated times, samples were denatured by SDS treatment, and resolved by agarose gel electrophoresis. (**F**) Loading reaction as in (E) after 60 min, using different combinations of ATP and MukBEF or the ATP-hydrolysis deficient E1407Q (EQ) mutant complex. (**G**) Loading reactions in the presence of topoisomerases. Reactions were performed as in (E), but an additional 50 mM NaCl was included in the reaction buffer, and DNA was nicked after BMOE treatment to adjust electrophoretic mobility. The experiment used pUC19 as the DNA substrate. (**H**) Loading on relaxed DNA substrates. DNA was relaxed by Topo I or nicking, purified, and used as in (E). Samples were nicked after BMOE treatment to make electrophoretic mobility comparable. The experiment used pUC19 as the DNA substrate. See also **Figure S1**, **Data S1**, and **Data S2**.

Although the Wadjet group covers a diverse sequence space, MukBEF has retained many key features of other SMC complexes ^2^ (**Figure 1B**). The SMC protein MukB dimerizes at its “hinge” domain, which connects via the long coiled-coil “arm” to the ABC-type ATPase “head” domain. MukBEF adopts a compact shape by folding over at its “elbow”, bringing the hinge close to the heads ^27–29^. The heads are bridged by the kleisin MukF, whereby the C-terminal winged-helix domain (cWHD) of MukF binds the “cap” surface of one MukB, and the N- terminal middle domain (MD) binds the coiled-coil “neck” of the other MukB. This designates the corresponding MukB subunits as κ- and ν-MukB, respectively. MukF also recruits the dimeric KITE protein MukE.

MukBEF is an obligate dimer, formed by two MukB_2_E_2_F monomers held together by their MDs and MukF N-terminal winged-helix domains (nWHD) ^18,30^. ATP binding induces engagement of the heads within a MukBEF monomer, enabling ATP hydrolysis and subsequent head disengagement ^18,31,32^. Cycles of head engagement and disengagement power the activities of all SMC complexes.

SMC complexes undergo turnover on DNA, with dedicated mechanisms mediating loading and unloading. This often involves loading factors such as Scc2/4, ParB, and Nse5/6, or unloading factors such as WAPL, MatP, XerD, and possibly microcephalin ^17,20,22,29,33–35^. Loading depends on ATP hydrolysis in MukBEF, cohesin, Smc–ScpAB, and Smc5/6, and involves the opening of a DNA entry gate, ingestion of DNA, and re-sealing of the gate ^17,18,20,29,36,37^. In principle, DNA entry can proceed via any of three candidate gates: the hinge gate, the neck gate, or the cap gate. Cohesin can load DNA through both its hinge and neck gates, whereas Smc5/6 loads through its neck gate exclusively ^20,33,36,38^. The neck gate also serves as cohesin’s exit gate for WAPL-mediated unloading ^39^. The entry and exit gates of MukBEF and other SMC complexes have not been identified.

DNA loading is complicated by the fact that MukBEF, and likely other SMC complexes, can entrap DNA as a “double-locked” loop with segments in separate compartments: the “ring” compartment, delineated by the kleisin, the SMC arms and the hinge, and the “clamp” compartment, delineated by the kleisin and the heads ^2,18–20^. In addition, entrapment of a single DNA segment in a post-extrusion “holding state” was recently observed for the MukBEF-related *E. coli* Wadjet I ^40^. The mechanistic basis for DNA transport into any of these compartments in any SMC complex is currently unclear. Here, we set out to investigate the loading process of MukBEF using biochemical reconstitution and cryo-EM reconstruction.

## Results

### Reconstitution of the MukBEF loading reaction

MukBEF loads onto chromosomal DNA to mediate long-range organization of the genome. We aimed to reconstitute DNA loading from purified components and enable its investigation by biochemical and structural methods. Previously, we monitored loading *in vivo* using site-specific covalent circularization of the MukB–MukF core by cysteine mutagenesis and BMOE-mediated cross-linking, inspired by work on cohesin and Smc–ScpAB ^15,17,18^. This strategy selectively probes for entrapment in the ring or a topologically equivalent compartment, and converts loaded complexes into SDS-resistant covalently closed protein-DNA catenanes. These can be separated from free or non-circularized complexes and detected by gel electrophoresis. We now adapted this assay from our *in vivo* setup to an *in vitro* setup using circular plasmids (**Figures 1C** and **S1B**). We employed *Photorhabdus thracensis* MukBEF, which is better behaved in cryo-EM experiments than its *E. coli* homologue, and engineered cysteine pairs for BMOE cross-linking into the *P. thracensis* proteins (**Figures S1C** and **S1D**). BMOE cross-linking of the purified complex produced a product pattern similar to what we previously observed for *E. coli* MukBEF ^18^ (**Figures 1D** and **S1E**). To verify whether the engineered complex was functional, we replaced the chromosomal *mukFEB* locus of *E. coli* with the *P. thracensis* version, with and without the cysteine substitutions, and including a HaloTag on MukB. The chimeric strains were viable on rich media at 37 °C (**Figure S1F**), indicating that *P. thracensis* MukBEF can substitute for its *E. coli* homologue and is at least partially functional even in the presence of the cysteine point mutations.

Next, we incubated the purified complex with negatively supercoiled plasmid DNA in a low-salt buffer containing ATP. At various timepoints of the reaction, we added BMOE to circularize the MukB–MukF core. Finally, we added buffer containing SDS to strip off complexes that were not catenated with the DNA, and resolved the products by agarose gel electrophoresis (**Figure 1E**). The assay produced a ladder of bands, with slower migrating species appearing as the reaction progressed. We interpret this as single plasmids catenated with one or more circularized protein complexes, where loading of multiple complexes becomes prevalent later in the reaction. DNA entrapment was not observed in the absence of ATP and was abolished when the ATP-hydrolysis deficient E1407Q (EQ) mutant of MukB was used (**Figure 1F**). These findings suggest that the reconstituted loading reaction strictly depends on ATP hydrolysis, reproducing a fundamental characteristic of MukBEF loading *in vivo* ^18^.

### DNA relaxation facilitates MukBEF loading

MukBEF directly binds topoisomerase IV (Topo IV) via its hinge region ^41^, and we wondered whether this enzyme may modulate MukBEF loading by changing the local geometry of the DNA. We tested loading of *P. thracensis* MukBEF in the presence of either *P. thracensis* Topo IV which decatenates and relaxes DNA, *E. coli* Topo I which relaxes DNA, or *E. coli* DNA gyrase, which supercoils its substrate rather than relaxing it. As before, we incubated MukBEF with negatively supercoiled plasmid in the presence of ATP with and without topoisomerase, but increased the salt concentration to support topoisomerase activity. Under these conditions, loading was less efficient, but still produced a distinctive ladder (**Figure 1G**). As a post-loading treatment after the addition of BMOE, we added a nicking enzyme to collapse DNA topoisomers and make the electrophoretic mobility of all samples comparable. We observed that loading was stimulated both by *P. thracensis* Topo IV and *E. coli* Topo I, but not by DNA gyrase (**Figure 1G**). Because Topo IV and Topo I relax DNA, but gyrase does not, we tested whether loading was also stimulated on relaxed substrates in the absence of the topoisomerase enzymes. We prepared DNA substrates relaxed either by Topo I treatment or by nicking, and subsequently purified the DNA. We then repeated the loading reaction under low-salt conditions in the absence of topoisomerases. After BMOE treatment, we again converted DNA to nicked open circles to adjust their mobility, added loading buffer with SDS, and resolved the reaction products by agarose gel electrophoresis (**Figure 1H**). Both nicked and relaxed substrates showed a strong increase in loading efficiency compared to negatively supercoiled DNA. This suggests that loading of MukBEF is influenced by the DNA topology, with a preference for environments where the DNA is less supercoiled or torsionally strained.

### ATP binding triggers opening of the neck gate

To gain detailed insights into the DNA transactions of MukBEF, we vitrified samples of the reconstituted loading reaction and analyzed them by cryo-EM. In addition to a sample under ATP turnover conditions, we collected datasets where sodium vanadate or beryllium fluoride had been added one hour after reaction start to enrich for species with engaged ATPase heads. All three conditions enabled the reconstruction of a state with engaged heads, and we pooled the datasets to increase the signal and obtain higher resolution (**Figure 2A** and Methods). The resolved state was free of DNA, and the neck gate had opened widely. We refer to this state as the “open-gate state” (PDB: 9GM7).

**Figure 2.**
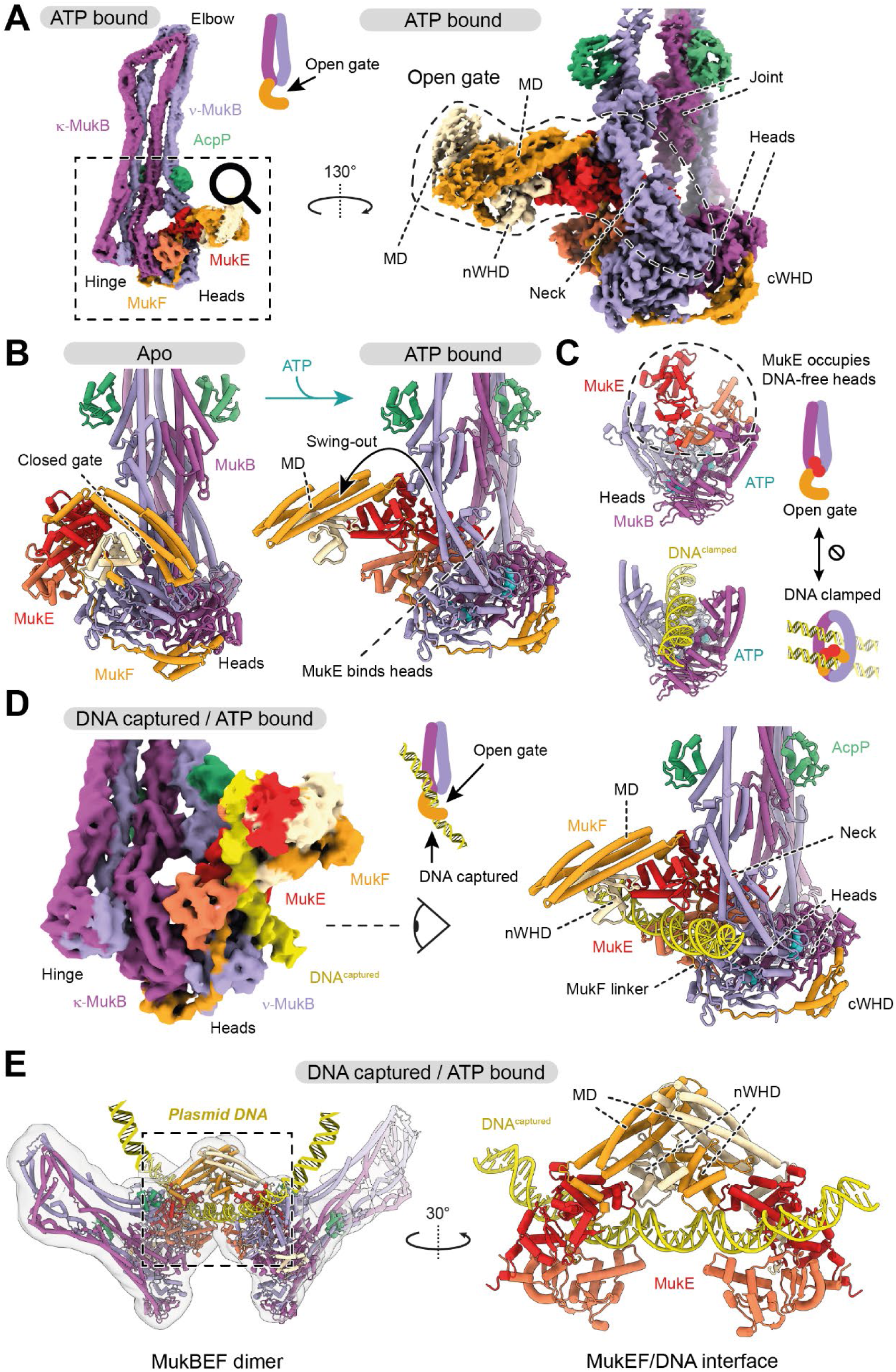
Mechanism of gate opening and DNA capture. (**A**) Structure of the open-gate state. Cryo-EM density of the MukBEF monomer in the nucleotide-bound form (left; PDB: 9GM7) and a focused refinement of the head module with open neck gate (right; PDB: 9GM8). (**B**) Comparison of apo (left; PDB: 7NYY)^18^ and open-gate state (right; PDB: 9GM8). Heads engage upon nucleotide binding, resulting in a swing-out of the MukF MD. (**C**) Comparison of the engaged MukB heads in the open-gate state (top; PDB: 9GM8) and the DNA-clamped unloading state (bottom; PDB: 7NYW)^18^. Binding of MukE and DNA to the top of the heads is mutually exclusive. (**D**) Structure of the DNA capture state. Focused classification of (A) reveals DNA captured at the open gate. Cryo-EM density (left) and cartoon model (right; PDB: 9GM9) are shown. (**E**) The DNA capture state in the context of the MukBEF dimer. Cryo-EM density of the dimer (blurred with a σ = 22 Å Gaussian filter to make low-density regions interpretable), cartoon model representation (left; PDB: 9GMA), and close-up of the dimeric DNA-capture interface (right) are shown. See also **Figure S2**.

The neck and head regions of MukB adopted radically different conformations from what we had previously observed for the apo and DNA-bound unloading states of MukBEF (**Figure S2A**). ATP binding and head engagement had triggered the detachment of the MukF MD from the MukB neck, which resulted in a swing-out of the MD of about 180° (**Figures 2B** and **S2B**). Detachment of the MD was facilitated by the mechanical distortion of the MukB neck constrained between engaged heads and aligned arms (**Figure S2A**), while the MD swing-out was stabilized by the binding of MukE to the top surface of the heads (**Figure 2C**). This surface is formed by the engagement of the heads upon ATP binding and is a highly conserved DNA binding site in all SMC complexes. Our structure reveals that occupation of the top of the heads by MukE and DNA is mutually exclusive, suggesting that MukE senses the DNA-free state of the heads to open the neck gate.

### DNA capture at the open neck gate

Focused sub-classification of the particle images revealed a DNA-bound structure (**Figure 2D**). The DNA was captured directly at the open gate, while the proteins adopted a conformation virtually identical to the open-gate state (**Figure S2C**). We refer to this structure as the “DNA capture state” (PDB: 9GM8). A low-resolution reconstruction of the dimeric MukBEF assembly in the capture state showed that both monomers bound a continuous DNA segment of about 52 bp (PDB: 9GMA; **Figure 2E**). DNA-binding surfaces were largely contributed by MukE and MukF, and to a lesser extent by the root of the ν-MukB neck. Compared to the apo state, MukE and MukF had aligned their DNA-binding surfaces to enable DNA capture (**Figure S2D**). The DNA was not entrapped inside the complex, but bound at its periphery without contacting the top-surface of the heads (**Figure S2E**). MukE employed a DNA binding mode overall similar to its role in the DNA clamping; however, the DNA followed a differently bent path along its surface (**Figure S2F**). As the captured DNA is positioned at the open neck gate but not entrapped, we reason that entrapment may be achieved by ingestion through the gate.

### Discovery of a bacteriophage MukBEF inhibitor

Is DNA capture involved in loading of MukBEF? A serendipitous discovery from phage biology helped us address this question, as will be explained in the following paragraphs.

Bacteriophage T7 infects *E. coli* and encodes the RecBCD inhibitor gp5.9, which interferes with the processing of DNA ends ^42–44^. Although RecBCD is not essential for host survival, we noticed that the production of gp5.9 from an arabinose-inducible promoter was highly toxic (**Figure 3A**). This was also the case in a Δ*recB* strain (**Figure 3B**), suggesting that the toxicity was not caused by a gain of function of gp5.9-bound RecBCD, but rather by targeting of another unknown and essential factor. To identify this factor, we immunoprecipitated FLAG-tagged gp5.9 (gp5.9^FLAG^) from wild-type (WT) and Δ*recB* extracts and analyzed the samples by TMT-MS (**Figures 3C**, **S3A**, and **S3B**). Both MukE and MukF were among the top hits, providing a possible explanation for the strong growth defect upon gp5.9 induction.

**Figure 3.**
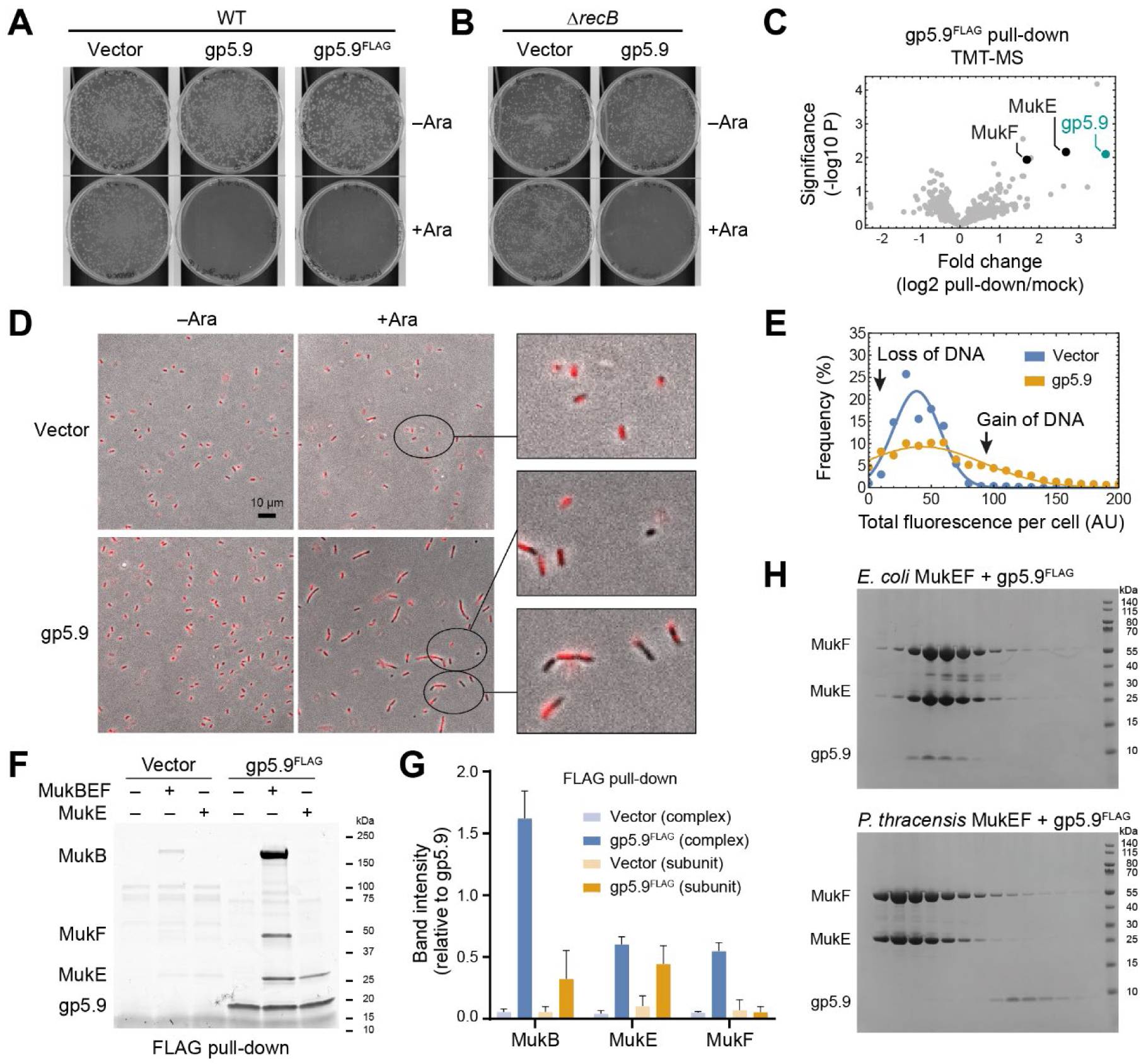
Discovery of a viral MukBEF inhibitor. (**A**) Expression of gp5.9 is toxic. *E. coli* cells were transformed with a kanamycin-selectable empty vector control or an equivalent construct containing gp5.9 under an arabinose-inducible promoter. Transformation reactions were plated on LB plus kanamycin with or without arabinose. Plates were incubated at 37 °C. (**B**) As in (A), but using a Δ*recB* background. (**C**) TMT-MS analysis of a gp5.9^FLAG^ pull-downs using pooled signal from WT and Δ*recB* extracts. A volcano plot of significance versus pull-down over mock extract is shown, highlighting gp5.9, MukE and MukF levels. (**D**) Morphology of cells expressing gp5.9. Cells carrying the indicated constructs were grown for three hours in LB media with or without arabinose, fixed with formaldehyde, stained with DAPI and imaged by combined phase contrast (grayscale) and fluorescence (red) microscopy. (**E**) Analysis of the DAPI intensity distribution of cells from the experiment shown in (D). Expression of gp5.9 causes a relative increase in cells with altered DNA content. (**F**) Pull-down of recombinant MukBEF or MukE with gp5.9^FLAG^. Anti-FLAG beads were charged with extract containing or lacking gp5.9^FLAG^, then incubated with recombinant MukBEF proteins, eluted with FLAG peptide and analyzed by SDS-PAGE and Coomassie staining. (**G**) Quantification of pull-downs as in (F), normalizing the indicated band intensities for the corresponding gp5.9^FLAG^ signal. Band intensities for MukB, MukE, and MukF are shown, comparing the signal between MukBEF complex and single subunit pull-downs. (**H**) SEC analysis of mixtures of gp5.9 and *E. coli* MukEF (top) and *P. thracensis* MukEF (bottom), respectively. Elution fractions were analyzed by SDS-PAGE and Coomassie staining. gp5.9 forms a stable complex with *E. coli* MukEF, but not with *P. thracensis* MukEF. See also **Figure S3** and **Data S3**.

Prompted by these findings, we investigated whether induction of gp5.9 caused chromosome segregation defects, a hallmark phenotype of cells with inactive MukBEF. Cells expressing gp5.9 produced more anucleate progeny than the empty vector control, which coincided with a higher fraction of cells with an increased DNA content (**Figures 3D, 3E**, **S3C**, and **S3D**). Cell width was unaffected by gp5.9 expression, whereas cell length was increased (**Figure S3E**), and many cells showed evidence of defective chromosome partitioning (**Figure S3C**). These findings suggest that gp5.9 interferes with chromosome segregation, consistent with the notion that it inactivates MukBEF.

Next, we investigated whether gp5.9 binds MukBEF directly. Recombinant MukE, MukEF, and MukBEF were efficiently pulled down by gp5.9^FLAG^-bound beads, whereas binding of MukB and MukF was lower or nearly undetectable, respectively (**Figures 3F**, **3G**, **S3F**, and **S3G**). This suggests that gp5.9 binds MukBEF mainly through the MukE subunit. In size exclusion chromatography (SEC), *E. coli* MukEF and gp5.9 formed a stable complex, whereas little if any binding was observed with *P. thracensis* MukEF (**Figure 3H**). Consistently, the *E. coli* strain with its endogenous *mukFEB* operon replaced by the *P. thracensis* version showed reduced susceptibility to gp5.9 (**Figure S3H**). As *E. coli* is the natural host for bacteriophage T7, these findings suggests that gp5.9 has evolved specificity for its target.

### gp5.9 targets the MukE DNA-binding cleft and inhibits DNA loading

To gain insights into how gp5.9 binds and inactivates MukBEF, we solved the structure of gp5.9 bound to *E. coli* MukEF by cryo-EM (PDB: 9GMD; **Figure 4A**). Focused classification, signal subtraction of the MukF core, and focused refinement using neural-network-based regularization with Blush ^45^ resolved a 73 kDa region of MukE bound to gp5.9. As observed previously in its RecBCD-bound form, gp5.9 formed a parallel coiled-coil dimer complemented by a beta-sheet of the N-terminal strands ^42^. gp5.9 bound along the DNA-binding cleft of MukE, overlapping along its full length with the DNA capture site (**Figure 4B**). Contacts of gp5.9 with MukE differed from those with RecBCD (**Figures S4A** and **S4B**), as did the precise path of DNA in the respective DNA binding sites (**Figure S4C**). However, the orientation of gp5.9 with respect to the DNA molecule is broadly similar, in the sense that the long axis of the gp5.9 coiled coil aligns approximately with that of the double helix. Moreover, within the resolution limits of the structures, gp5.9 positioned several negatively charged residues (D11, D15, D21, E24, E36, D38, E43, E45) near positively charged residues in MukE (R140, K150, R163, R164, R179) or MukF (R322). Most of these ion pair interactions mimic DNA phosphate backbone contacts and thus prevent the natural DNA substrate from binding efficiently (**Figure S4D**). Therefore, although there are significant differences in the details of binding to individual targets, the data overall support the designation of gp5.9 as a DNA mimic protein.

**Figure 4.**
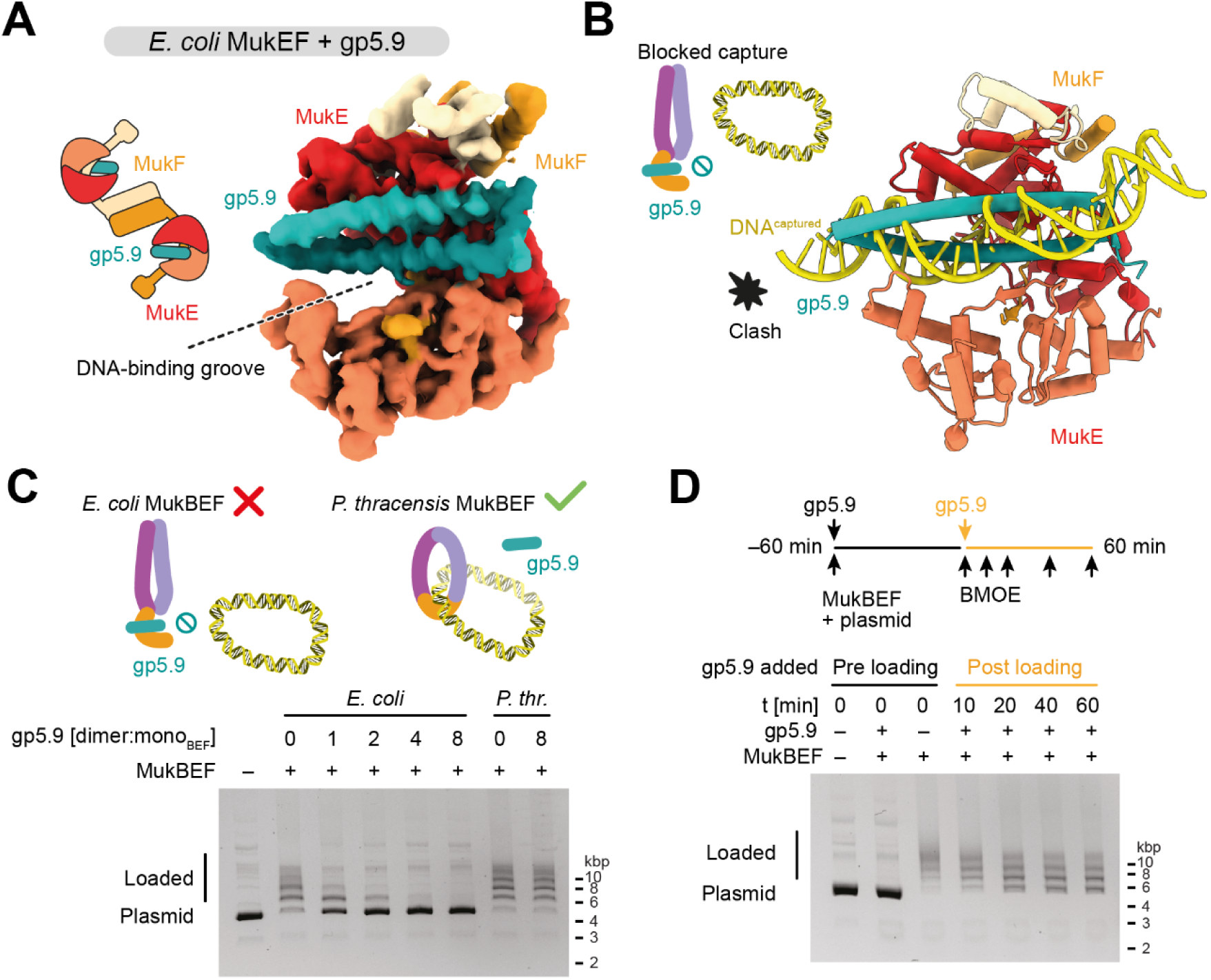
gp5.9 binds the DNA capture site and inhibits loading. (**A**) Structure of the gp5.9/MukEF interface. A cartoon of the complex analyzed (left) and cryo-EM density from a focused refinement (right) is shown. (**B**) DNA capture and gp5.9 binding are mutually exclusive. The cartoon representation of (A) is shown (PDB: 9GMD) with DNA from the superimposed capture state structure (PDB: 9GM9). (**C**) DNA entrapment assay in the presence of gp5.9 as in Figure 1H using nicked plasmid (pUC19). The molar ratio of gp5.9 to MukBEF monomer sites is indicated. *E. coli* MukBEF is sensitive to gp5.9, whereas *P. thracensis* MukBEF is not. (**D**) As in C, but gp5.9 was added 60 min after reaction start. Samples were then treated with BMOE at the indicated timepoints after addition of gp5.9. See also **Figure S4**.

The structure revealed that binding of gp5.9 to the MukE DNA-binding cleft is mutually exclusive with formation of the DNA capture state. Therefore, if the capture state does indeed take part in DNA loading, then gp5.9 would be expected to inhibit the loading reaction. To test this, we prepared purified *E. coli* MukBEF containing cysteine pairs for covalent circularization. Like the *P. thracensis* complex, *E. coli* MukBEF efficiently produced an SDS-resistant ladder of plasmid-bound species after loading and BMOE cross-linking (**Figure 4C**). When gp5.9 was added to the reaction, we observed a strong inhibition of ladder formation, with an almost complete loss at a two-fold molar excess of gp5.9 over MukBEF. In contrast, loading of *P. thracensis* MukBEF was unaffected even by an 8-fold molar excess, highlighting again the specificity of gp5.9 inhibition.

We reasoned that the effect of gp5.9 on MukBEF loading may be explained by two scenarios: an inhibition of loading or, alternatively, an acceleration of unloading. To dissect its mode of action, we performed the following experiments. Addition of an 8-fold molar excess of gp5.9 at different timepoints quenched the loading reaction at intermediate levels of DNA entrapment (**Figure S4E**). When loading reactions were run for one hour, then quenched with gp5.9 and incubated for an additional hour in the presence of the inhibitor, only modest unloading was observed (**Figure 4D**). This effect, if caused by gp5.9 at all, cannot explain the strong entrapment defect observed when gp5.9 was included at early timepoints of the reaction. In summary, these results suggest that gp5.9 inhibits DNA loading and support the notion that DNA capture is necessary for DNA entrapment.

## Discussion

### Neck gate opening in SMC complexes

The entrapment of DNA by SMC complexes requires the passage of DNA through an entry gate. Our findings show that MukBEF employs a dedicated mechanism for opening its neck gate, converting the DNA-free apo form to the open-gate state: 1) ATP binds the heads and leads to their engagement, 2) the neck distorts and releases the MD of MukF, 3) MukE binds the DNA-free top of the heads and stabilizes the open-gate state. This mechanism ensures that the gate opens only when the heads are DNA-free, which serves as an indicator that the complex is ready for loading. In line with this idea it has been found that ATP-induced neck gate opening in condensin and cohesin can be suppressed by linear double-stranded DNA ^29,33,46,47^. This suggests that these complexes may employ a selective gating mechanism similar to that of MukBEF. Opening of the neck gate in Smc5/6, in contrast, differs from the mechanisms used by cohesin, condensin and MukBEF, as it only requires Nse5/6 but not ATP ^20,48,49^. Although it may be controlled in distinct ways, neck gate opening emerges as a central property of SMC complexes.

### The DNA capture state as a first step of loading

DNA entry into an SMC complex works against a large entropic cost, making it more likely for DNA to be positioned outside than inside, and rendering stochastic gate passage inefficient. Analogous to the directed transport of molecules across biological membranes an initial substrate capture step may help to guide DNA through the entry gate. We propose that the DNA-bound structure obtained here represents this capture state.

Is the DNA capture state involved in DNA loading? We find that gp5.9 targets the DNA-binding site of MukE, which contacts DNA both in the capture state and when DNA is entrapped in the clamp compartment. As gp5.9 inhibits the loading reaction, either form of DNA binding may be involved in loading. We favor the capture state as the relevant target for the following reasons: DNA entrapment in the clamp requires ATP hydrolysis *in vivo*, indicating that it occurs after loading ^18^. Structural evidence and *in vivo* entrapment assays also suggest that DNA entrapment in the clamp coincides with entrapment in the ring compartment, implying that clamping is a result, and not a precursor, of DNA loading ^18^. The capture state, however, requires ATP binding only but not hydrolysis, and can thus occur before DNA entrapment. This makes it an attractive first step of the loading reaction.

### Mechanism of DNA entry through the neck gate

Combining our new structures with existing data now enables us to propose a pathway of DNA loading through the neck gate (**Movie S1**). This mechanism only requires a single round of ATP hydrolysis, which will be explained in the following. A recent structure of another member of the Wadjet family, *E. coli* Wadjet I, was solved in a post-hydrolysis state after DNA loading and loop extrusion, called the “holding state” ^40^. The holding state entraps DNA in a compartment formed by the kleisin JetA/MksF and the head-proximal part of JetC/MksB. Comparison with the MukBEF capture state suggests a straightforward conversion reaction (**Figures 5A-C**, and **S5A**). Starting with the ATP- and DNA-bound capture state, we envision that upon ATP hydrolysis the MukB subunits revert to their apo conformation. This has two major conformational consequences: 1) disengagement of the composite surface on top of the heads, and 2) straightening of the MukB neck. As both transitions are incompatible with binding of MukE to the MukB heads, MukB will release from MukE and the DNA. However, MukB cannot diffuse away because it is tethered to MukF via the cWHD and flexible linker (**Figures 5C** and **S5A**). MukB is free to sample the space around the DNA, and as its straightened neck is now competent to bind the MD of MukF, the neck gate will eventually close. This results in an overall rotation of MukB that wraps MukF around the DNA and generates the holding state with DNA entrapped inside (**Figures 5B** and **5C**).

**Figure 5.**
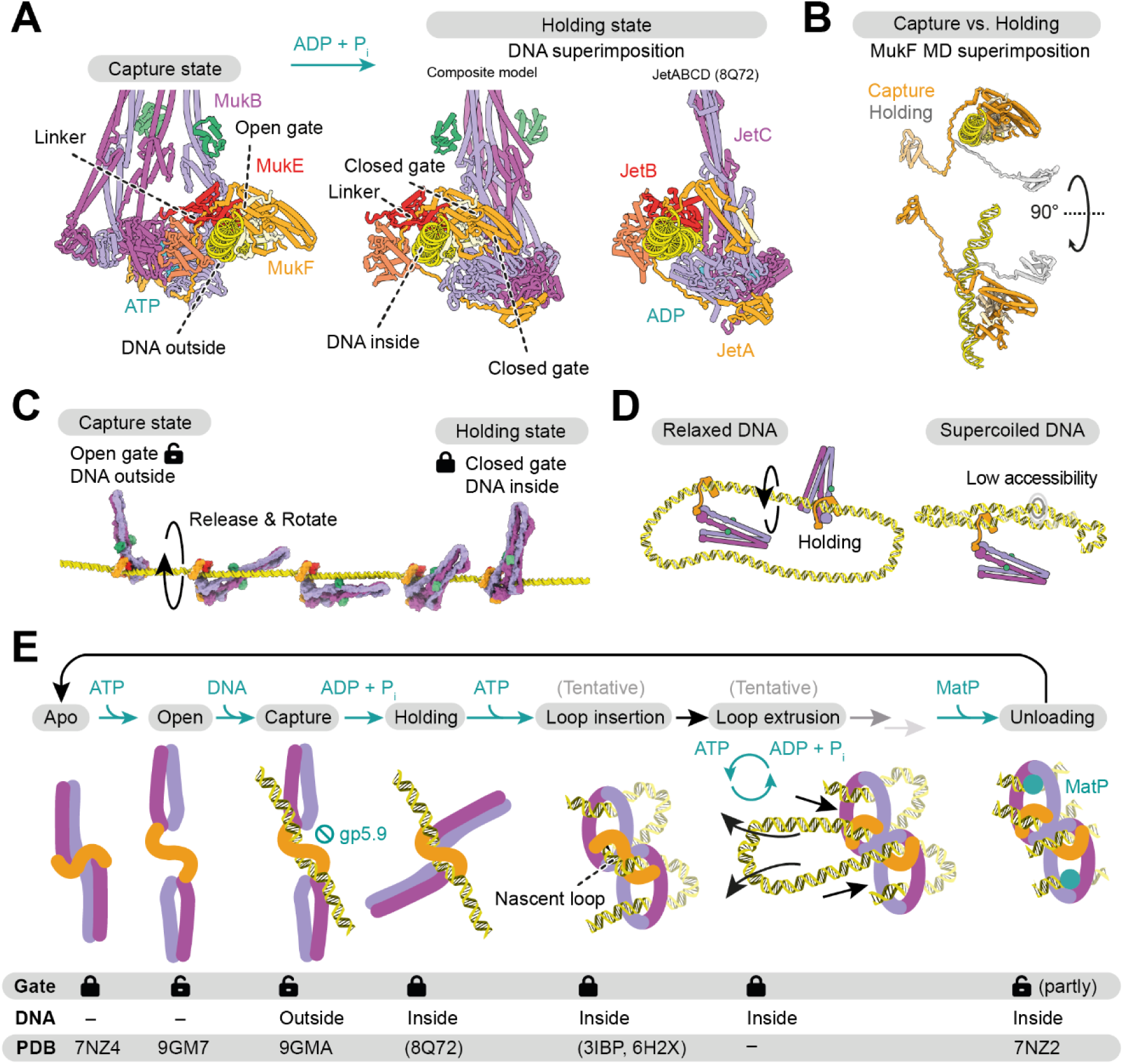
Mechanism of DNA entry into MukBEF. (**A**) Comparison of the DNA capture state (left) with the *E. coli* Wadjet I holding state (right; PDB: 8Q72)^40^, and a model of the equivalent MukBEF holding state (middle). The latter was composed from DNA-bound MukEF (PDB: 9GM9), the apo MukB/MukF interface (PDB: 7NYY)^18^, and a remodeled MukF linker. Coordinates were superimposed on the DNA. The state transition from capture to holding state requires a rotation of MukB and the MukF linker around the DNA. (**B**) Comparison of MukF between capture and holding state. The linker wraps around DNA upon the proposed state transition. (**C**) “Release & rotate” model for transition from the capture to the holding state and gate closure. MukB releases from MukE upon ATP hydrolysis and rotates around the DNA to close the neck gate. (D) Implications of the release & rotate model for loading on relaxed (left) and supercoiled (right) DNA. Rotation around a relaxed double-strand is easier than in the context of a compact plectoneme, and is consistent with the inhibition of loading on supercoiled DNA. (E) Model of the MukBEF activity cycle. The state of the neck gate and entrapment of DNA are indicated, and PDB IDs that support the states are shown. Parentheses around IDs indicate partial or homologous structures. Three-dimensional models for the tentative states are available in **Data S4.** See also **Figure S5**, **Data S4** and **Movie S1**.

This “release and rotate” model of DNA entrapment has several attractive properties. First, the model explains how DNA loading depends on ATP hydrolysis. While ATP binding exposes the DNA capture site, ATP hydrolysis triggers closing of the neck gate and ingestion of the captured DNA. Second, the model explains why loading is more efficient on relaxed DNA and may benefit from a cooperation with topoisomerases: Rotation of MukB around the DNA needs space, and relaxation makes the double strand more accessible compared to a plectonemal supercoil (**Figure 5D**). Notably, folding at the elbow reduces MukB’s radius of gyration, which may facilitate this movement. Third, the product of the loading reaction, the holding state, is consistent with our entrapment assay, which converts it into a protein/DNA catenane. Finally, the loading model predicts the start site of DNA loop extrusion. Transition from the capture state to the holding state retains a short DNA segment at the center of the MukBEF dimer. This segment is equivalent to the extruded loop in the *E. coli* Wadjet I post-extrusion holding state ^40^ (**Figure S5B**). Our model thus predicts that extrusion initiates directly at the captured DNA segment.

### Switching from DNA loading to DNA loop extrusion

Both DNA loading and DNA loop extrusion require ATP hydrolysis ^17,18,50–52^. We propose that these processes are separate and use the ATPase cycle in different modes. While gate opening is a prerequisite for loading, it is likely detrimental to loop extrusion and needs to be suppressed during the operation of the motor. Our findings suggest how this is achieved, and how the switch from “loading mode” to “loop extrusion mode” may be implemented: Once DNA is inserted into the clamp during extrusion, the top surface of the heads becomes inaccessible to MukE, blocking the gate opening mechanism described above.

How can MukBEF insert DNA into the clamp and switch to loop extrusion? Starting from the holding state, the clamped conformation can be generated by head engagement and tilting of the MukEF-bound DNA segment onto the top of the heads (**Figures 5E** and **S5C**, and **Movie S1**). This results in the overall insertion of a DNA loop, which is “double-locked” in ring and clamp compartments, as supported by the structure of the MatP-bound unloading state and cross-linking studies. We envision that the double-locked loop is part of the extrusion reaction, as proposed previously ^18^. Consistent with this notion, cross-linking experiments with condensin and Smc5/6 suggest that these complexes also insert double-locked loops^19,20^.

Although the exact mechanism of loop extrusion is unknown, it is conceivable that it involves the opening of the SMC arms. Structures of the MukB elbow in an extended conformation and the MukB hinge in an open V-shaped conformation support this idea ^27,53^ (**Figure S5C** and **Movie S1**).

In summary, we propose that a single ATP binding and hydrolysis cycle mediates the loading of MukBEF. The loading mode is specifically activated in DNA-free MukBEF, and once loaded, MukBEF can insert DNA into the clamp. This switches the complex to loop extrusion mode by suppressing further gate opening events, which may then become dependent on specialized unloading factors such as MatP.

### Inhibition of SMC complexes by pathogens

Several SMC complexes contribute to the defense against pathogens: Smc5/6 silences transcription of some viral genomes, cohesin participates in the recombination of immunoglobulin loci, and many members of the Wadjet group clear plasmid infections by specific activation of a nuclease ^6,12,40,54^. It is not surprising that pathogens have developed strategies to interfere with some of these processes: The Hepatitis B protein X (HbX) flags Smc5/6 for degradation, and the HIV-1 protein Vpr mediates the degradation of the Smc5/6 localization factor SLF2 ^12,54^. Here, we describe an inhibitory mechanism orthogonal to protein degradation: the blocking of DNA loading by the bacteriophage protein gp5.9.

Bacteriophage T7 encodes several inhibitors that inactivate host defenses or housekeeping functions, such as Ocr, which inhibits restriction enzymes, the BREX defense system and the host RNA polymerase. Furthermore, gp2 also inhibits the host RNA polymerase, gp0.4 inhibits the cell division protein FtsZ, and gp5.9 inactivates the RecBCD nuclease involved in recombination and degradation of linear DNA ^42,55–59^. gp5.9 is an acidic protein and considered a DNA mimic ^60,61^. We show here that it inhibits *E. coli* MukBEF but not *P. thracensis* MukBEF, and that its binding mode to MukBEF is different from its binding to RecBCD. Although gp5.9 targets DNA binding sites by contacting residues involved in phosphate backbone binding, it encodes sufficient specificity to interfere with select targets. This “tailored” mimicry is a common theme among the structurally diverse group of viral DNA mimics, such as anti-CRISPR and anti-restriction proteins ^60,61^.

Like several other members of the Wadjet group, MukBEF is lacking the MksG/JetD nuclease, and is unlikely to restrict pathogens by genome cleavage. It is currently unknown whether MukBEF protects against phage infection at all, or whether gp5.9 targets MukBEF as part of a more general assault against the host’s metabolism. Since gp5.9 function is not essential for T7 propagation ^44,62^, we suspect that MukBEF inhibition is required only under certain conditions, or for maintaining the long-term competitive fitness of the virus.

### Outlook

Our findings reveal a novel mechanism of SMC inhibition, and we anticipate that more anti-SMC proteins will be discovered in future studies. For example, MatP unloads MukBEF from chromosomes, and pathogens could potentially exploit related strategies to guard their genomes against SMC activity.

Gate opening and topological DNA entrapment are widely recognized as essential for sister chromatid cohesion, a specialized function unique to the cohesin complex. However, the involvement of gate opening and topological entrapment in DNA loop extrusion remains debated, possibly due to the necessity for indirect methodologies^19,63–68^. Here, we directly visualized gate opening in a bacterial SMC complex and identified a novel DNA capture step that positions DNA at the open gate. We suggest that the sequence of gate opening, DNA capture and DNA entrapment must be considered a universal mechanism underlying SMC function by loop extrusion.

The structural evidence presented here supports a robust model for how DNA entrapment is achieved. It is now critical to investigate the next steps in the reaction cycle, namely how DNA loop extrusion capitalizes on DNA entrapment and uses ATP hydrolysis to generate folded chromosomes.

### Limitations of the study

Our loading model invokes a pre-extrusion holding state, which is closely related to the post-extrusion holding state of *E. coli* Wadjet I but lacks an experimental structure. It is thus possible that the product of MukBEF loading deviates from what we propose. In addition, the structures presented here were obtained by single-particle methods involving stringent subset selection, and thus explain only a fraction of the data. Other states may exist that are more flexible and cannot be averaged, are rare, or were missed due to inadequate selection strategies. Our efforts have also not revealed if and how bacteriophage T7 benefits from the inhibition of MukBEF, which will be a subject of future studies.

## Acknowledgements

We thank Gemma Fisher (MRC LMS) for the gift of purified MukBEF proteins for preliminary pull-down studies; Dan Taylor for assistance with protein purification; Stephen Cross (University of Bristol Imaging Facility) for assistance with image analysis; Kate Heesom and Phil Lewis (University of Bristol Proteomics Facility) for assistance and advice with TMT analysis and interpretation; Giuseppe Cannone and all members of the LMB electron microscopy facility for excellent EM training and support; Toby Darling, Jake Grimmett and Ivan Clayson (LMB scientific computing) for computing support. We thank Madhu Srinivasan and Kim Nasmyth for helpful discussions. F.B. acknowledges support by an EMBO Advanced fellowship (ALTF 605-2019) and the Wellcome Trust (227260/Z/23/Z). M.D. acknowledges support by the BBSRC (BB/Y004426/1). This work was funded by the Medical Research Council as part of UKRI (U105184326 to J.L.).

## Author Contributions

F.B. performed protein purifications, DNA entrapment assays, cryo-EM sample preparation, cryo-EM data acquisition and analysis, model building, strain construction, and bioinformatic analysis; B.C. performed toxicity tests and TMT proteomics; S.K. performed *in vitro* pull-downs and light microscopy; O.J.W. performed protein purifications and contributed to pull-downs, proteomics, light microscopy, and toxicity tests; D.K. advised on cryo-EM analysis and performed refinement of the gp5.9/MukEF structure with Blush; M.S.D. and J.L. supervised the study; F.B. prepared the manuscript with contributions from all authors.

---

**Figure S1.**
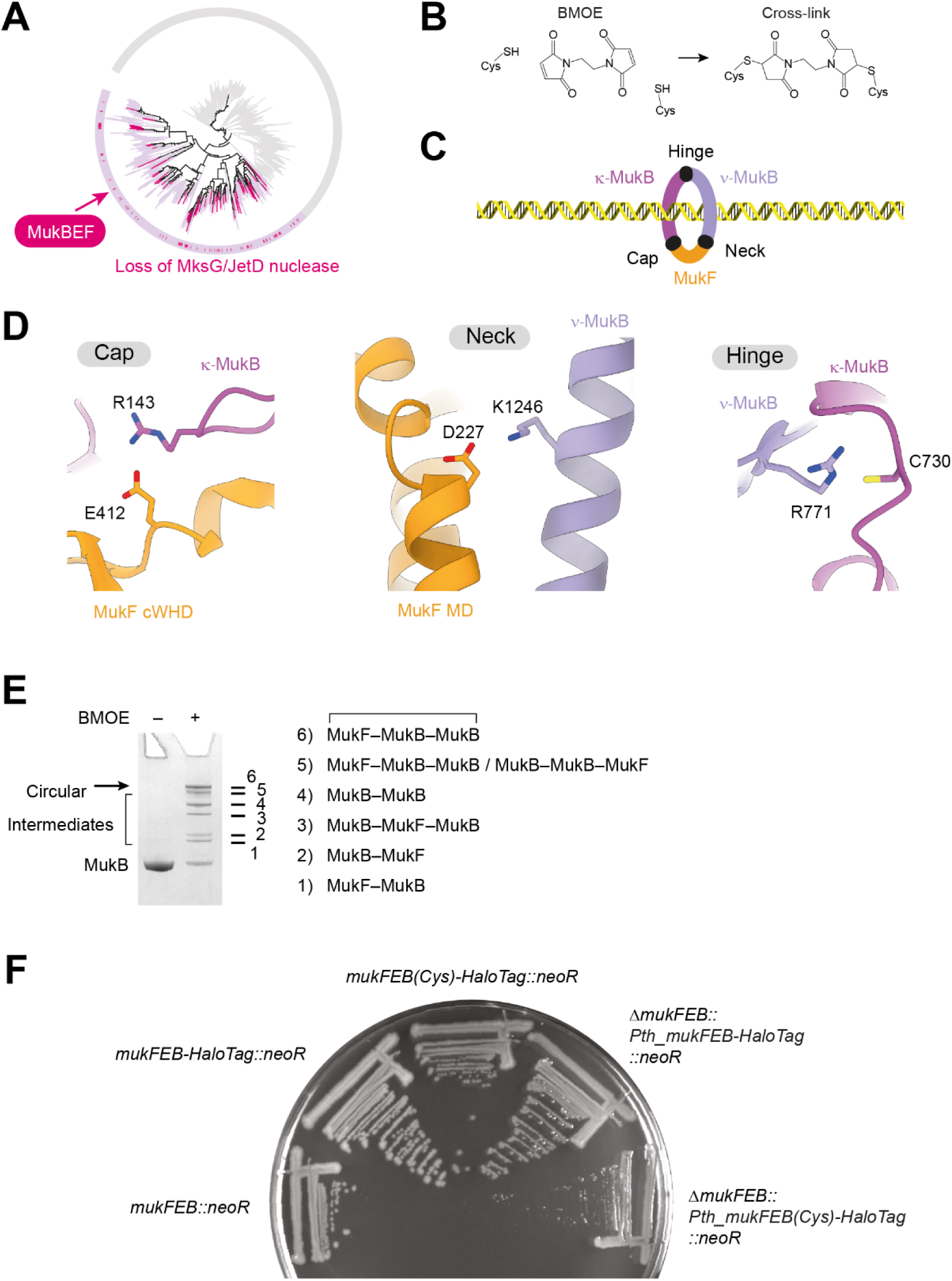
Phylogeny of MukBEF and cysteine mutagenesis. (**A**) Loss of the MksG/JetD nuclease across the Wadjet group. Absence of the nuclease gene is shown on the phylogenetic tree from **Figure 1A**. (**B**) BMOE cross-linking reaction between cysteine pairs. A covalent bridge between the cysteine sulfur atoms is formed. (**C**) Location of the three potential gates shown in the simplified cartoon representation of MukBEF. (**D**) Location of the residues in *P. thracensis* MukBEF targeted by cysteine mutagenesis. Residues are shown in the apo state (PDB: 7NYY). (**E**) Product assignment of the cross-linking reaction shown in **Figure 1D**, inferred from the closely related band pattern observed for *E. coli* MukBEF *in vivo* ^18^. (**F**) Growth of *E. coli* strains with the *P. thracensis mukFEB* locus substituted for the endogenous *mukFEB* locus. Strains were streaked for single colonies on TYE, and grown for 14 h at 37 °C. Note that the cysteine mutant *P. thracensis* variant causes a mild growth defect. Strains used: SFB012, SFB017, SFB174, SFB208, SFB209. See also **Figure 1**.

**Figure S2.**
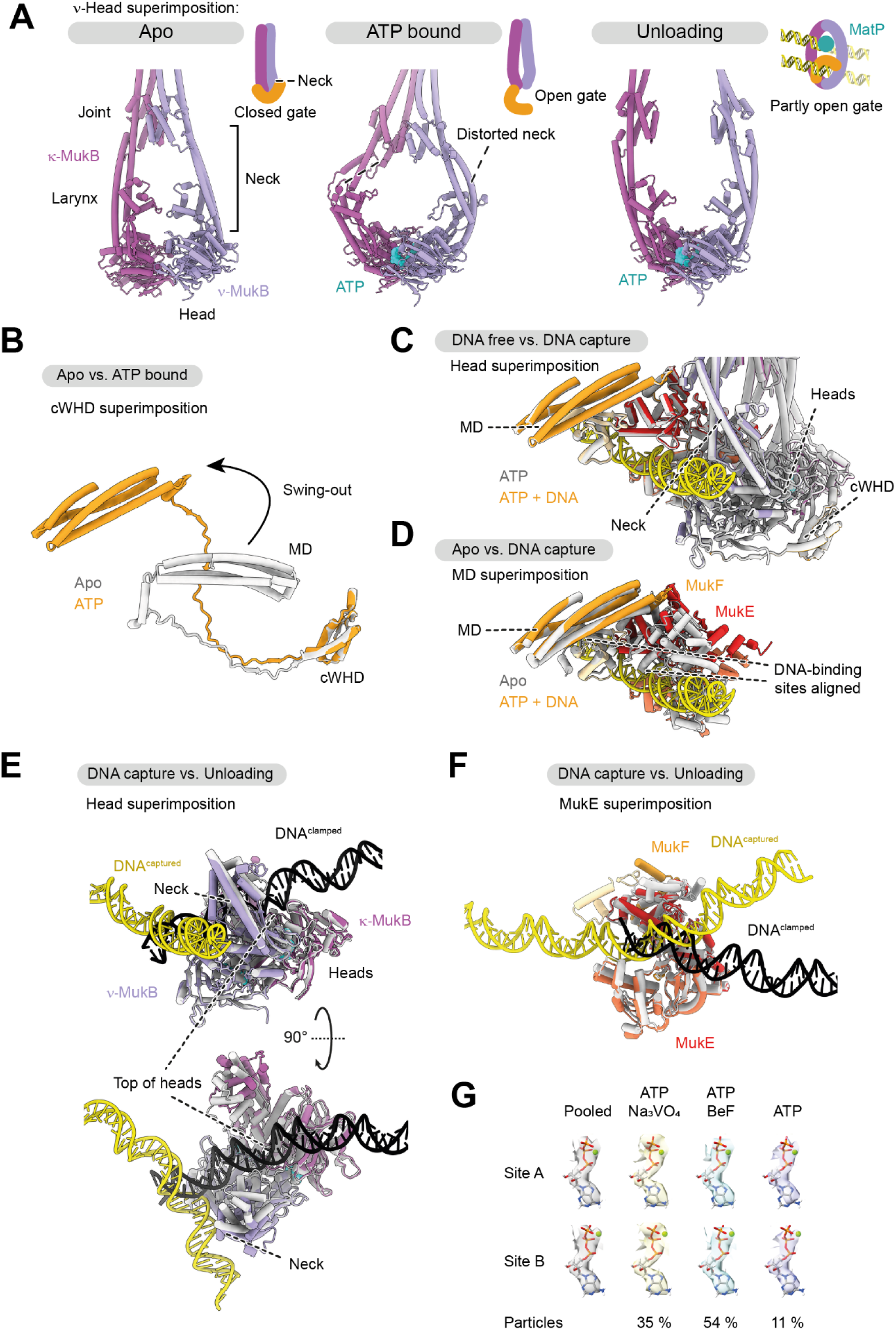
MukBEF conformations and DNA binding. (**A**) Conformations of the MukB head and neck region in the apo state (left; PDB: 7NYY), ATP-bound open-gate state (middle; PDB: 9GM6), and DNA-bound unloading state (right; PDB: 7NYW). The open-gate state has a severely distorted neck. (**B**) Comparison of MukF in apo state (gray; PDB: 7NYY) and ATP-bound open-gate state (colored; PDB: 9GM8). The MD swings out upon ATP binding. Structures were superimposed on the cWHD. (**C**) Comparison of the open-gate state (gray; PDB: 9GM8) and capture state (colored; PDB: 9GM9). Structures were superimposed on the heads. (**D**) Comparison of the apo state (gray; PDB: 7NYY) and capture state (colored; PDB: 9GM9). Structures were superimposed on the MD. The DNA-binding surfaces of MukE and MukF align in the capture state. (**E**) Comparison of DNA capture state (colored; PDB: 9GM8) and DNA unloading state (gray, black; PDB: 7NYW). Structures were superimposed on the heads. (**F**) Comparison of DNA binding to MukE in the DNA capture state (colored; PDB: 9GM9) and DNA unloading state (gray, black; PDB: 7NYW). Structures were superimposed on the MukE dimer. (**G**) Comparison of nucleotide cryo-EM density for reconstructions from individual datasets. The structure was refined against the pooled dataset, and individual maps were reconstructed using the particle poses obtained from this consensus refinement. Density in a zone of 2.5 Å around the nucleotide of both ATPase sites is shown, and the fraction of particles in the respective dataset is indicated. The nucleotide was modeled as MgATP. See also **Figure 2**.

**Figure S3.**
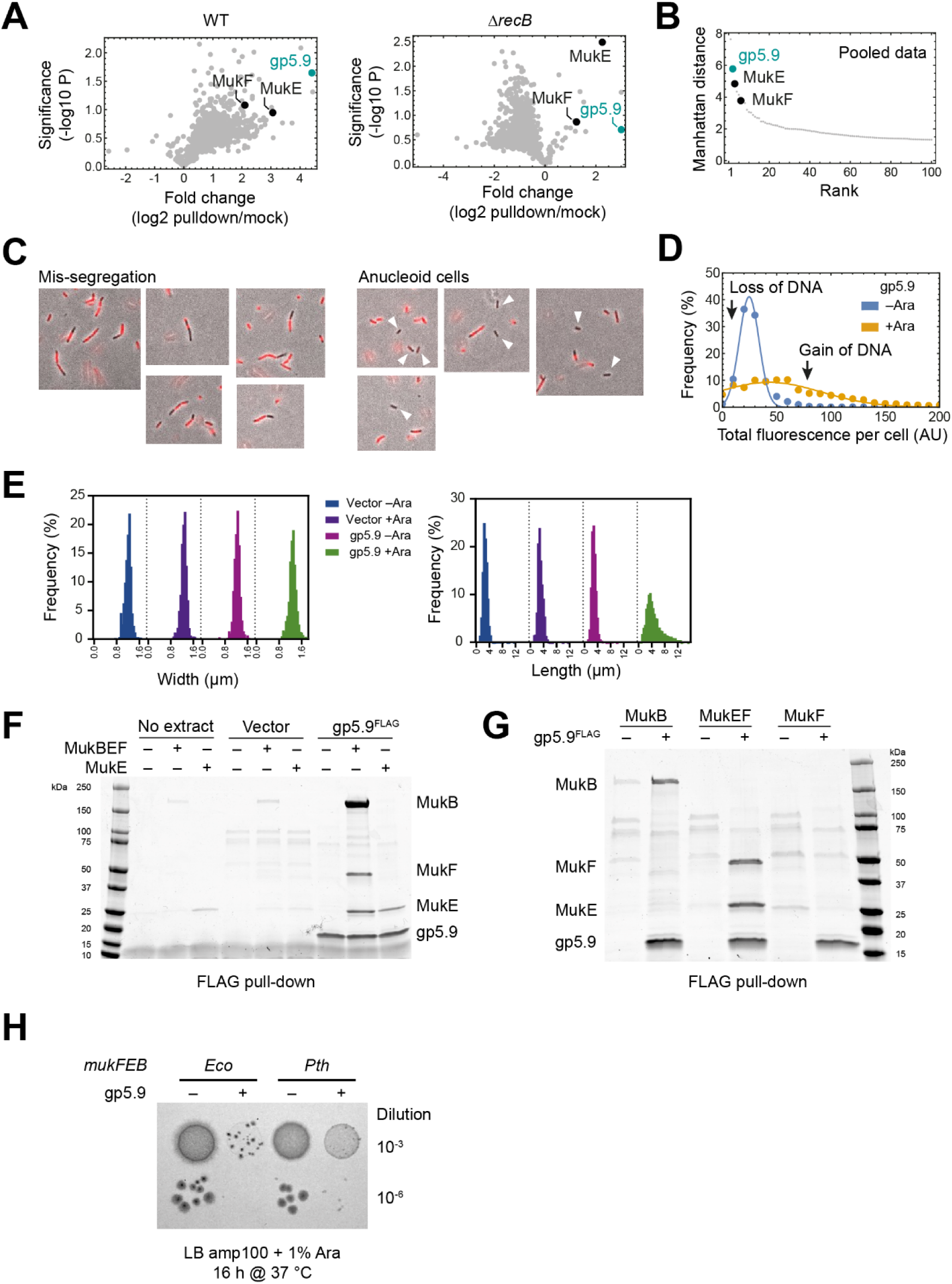
Effects of gp5.9 expression in E. coli. (**A**) TMT-MS analysis as in **Figure 3C**, showing unpooled data for WT and Δ*recB* extracts. (B) Manhattan distance ranking of the datapoints shown in **Figure 3C**. (**C**) Examples of chromosome mis-segregation and anucleate cell formation in cells expressing gp5.9. Anucleate cells are indicated by white triangles. (**D**) DAPI intensity distributions as in **Figure 3E**, comparing uninduced and induced conditions. (**E**) Cell width (left) and length (right) distributions of the experiment shown in **Figure 3D**. (**F**) Full gel shown in **Figure 3F**, also showing a pull-down of recombinant protein in the absence of extract. (**G**) Pull-down as in **Figure 3F**, using MukB, MukEF and MukF proteins. (**H**) gp5.9 sensitivity of *E. coli* with the endogenous *mukFEB* locus (*Eco*) replaced by the *P. thracensis* locus (*Pth*). Strains contained an ampicillin-selectable empty vector control or produced gp5.9 from an equivalent arabinose inducible construct. The indicated dilutions were spotted on LB media plus ampicillin with arabinose and incubated at 37°C. While the *Eco* strain only produced few colonies at the low dilution, the *Pth* strain produced a lawn at the same dilution, and single colonies at the low dilution. Note that the *Pth* strain still showed a growth phenotype upon gp5.9 induction, likely due to residual inhibition of MukBEF or inactivation of other targets such as RecBCD. Strains used: SFB289, SFB290, SFB292, SFB293. See also **Figure 3** and Data S3.

**Figure S4.**
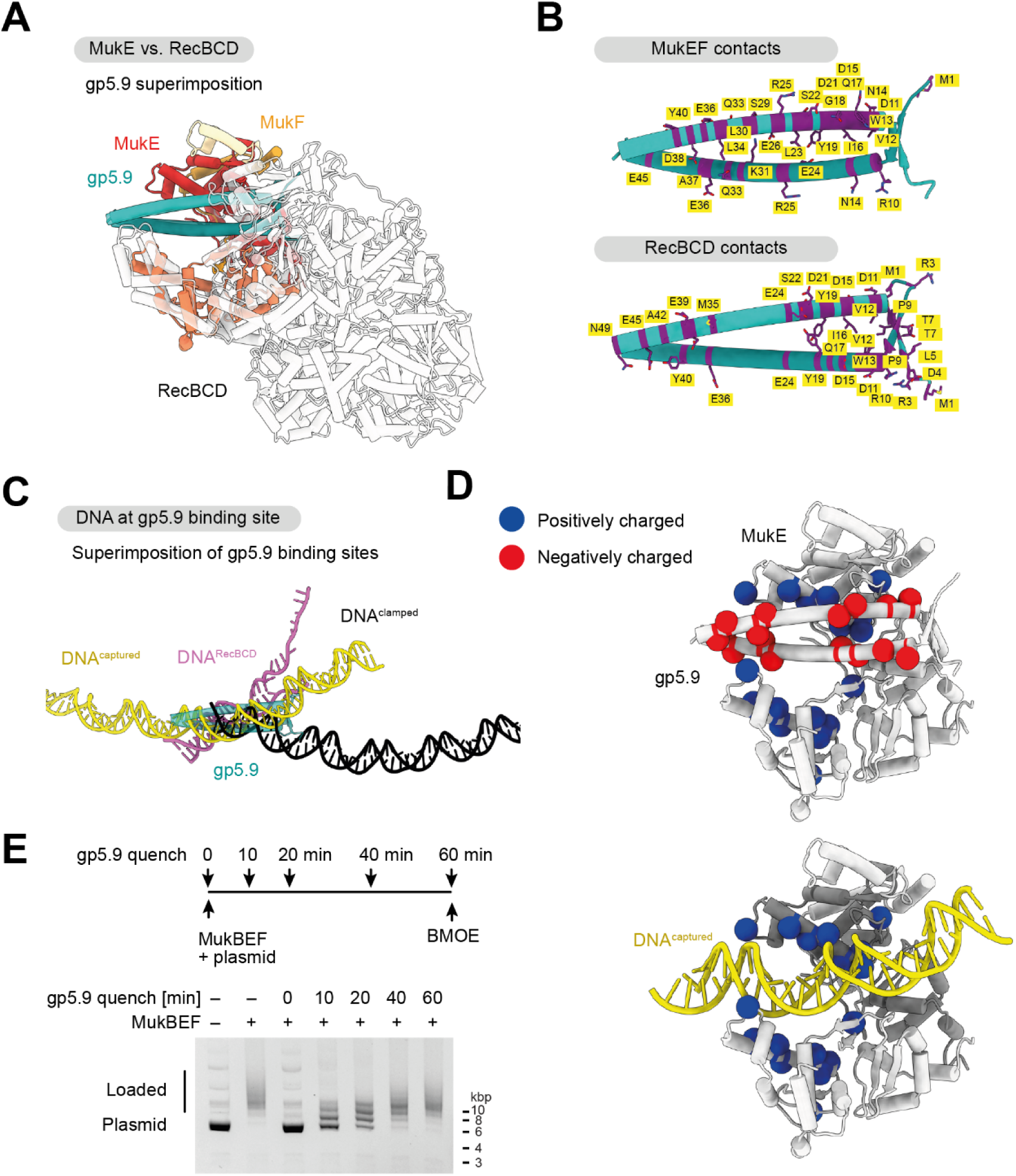
gp5.9 binding of RecBCD and MukBEF. (**A**) Comparison of gp5.9 binding to MukEF (PDB: 9GMD) and RecBCD (PDB: 8B1R)^42^. Structures were superimposed on gp5.9. (**B**) Binding residues on gp5.9 in the MukEF structure (top; PDB: 9GMD) and the RecBCD bound form (bottom, PDB: 8B1R). Residues with an inter-model atom-atom center distance of less or equal than 4 Å are highlighted in purple. (**C**) Comparison of DNA paths at the gp5.9 binding site. gp5.9-bound MukEF and RecBCD were superimposed on gp5.9 as in (A), and then DNA-bound forms were superimposed onto MukE or RecB, respectively. DNA paths are shown for the MukBEF DNA capture state (yellow; PDB: 9GM9), DNA-bound RecBCD (pink; PDB: 5LD2)^69^, and the MukBEF DNA unloading state (black; PDB: 7NYW)^18^. Superimposed gp5.9 (teal) are shown for reference. (**D**) gp5.9 places negatively charged residues (red) close to positively charged residues (blue) in the MukE DNA-binding cleft. C-alpha positions are shown as colored spheres (gp5.9: D11, D15, D21, E24, E36, D38, E43, E45; MukE: R140, K150, K154, R156, R161, R163, R164, R179; MukF: R322). The position of DNA in the capture state is shown for reference (bottom; structures superimposed on MukE). (**E**) Quenching of DNA loading by gp5.9. Entrapment assay as in **Figure 4C**, but an 8-fold molar excess of gp5.9 was added at the indicated timepoints. All samples were BMOE treated 60 min after reaction start. See also **Figure 4**.

**Figure S5.**
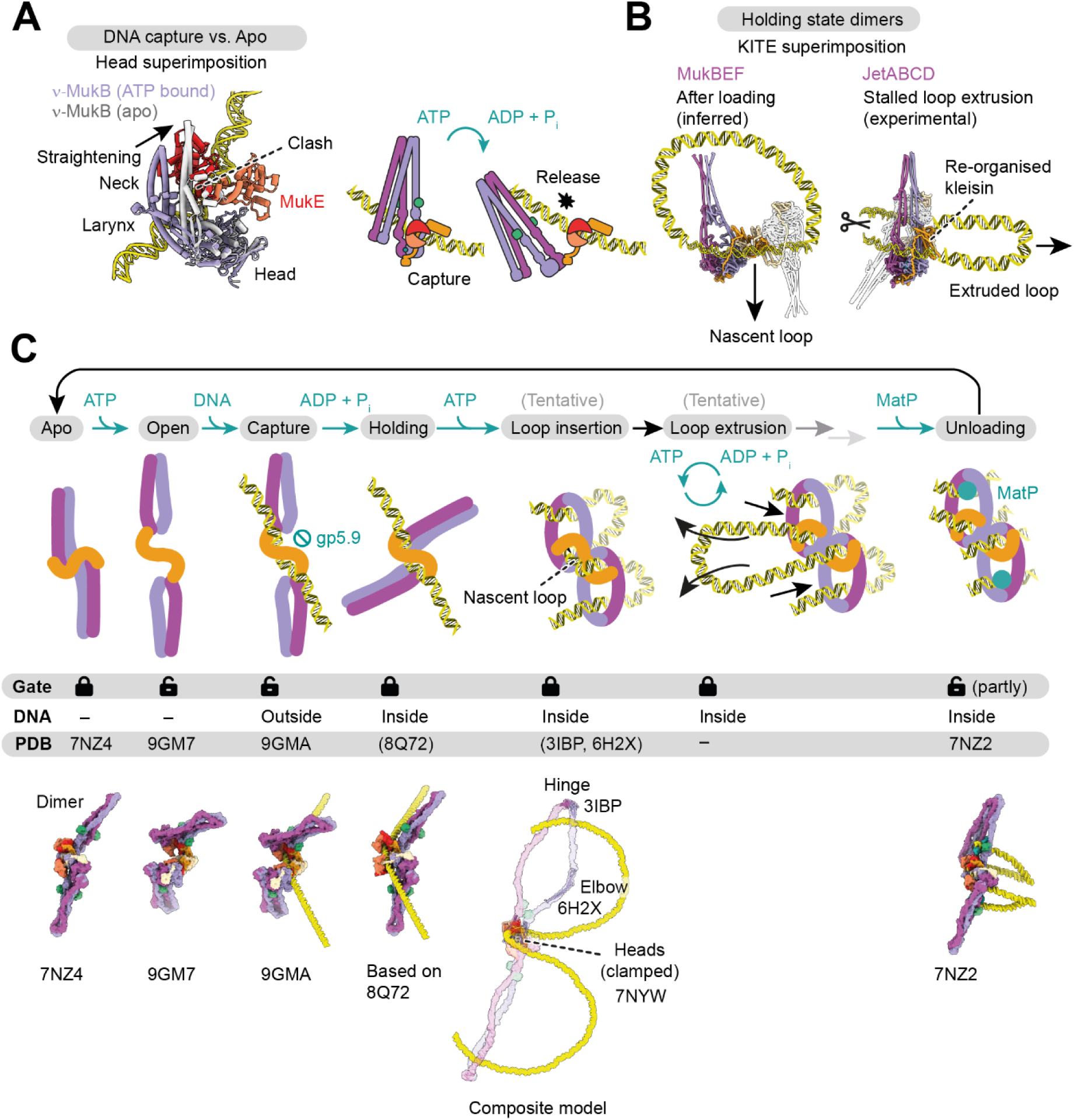
Structural support for DNA loading and loop extrusion. (**A**) Release of MukB upon ATP hydrolysis. A comparison of the DNA capture state (colored) with MukB in the apo state (gray) is shown on the left. Structures were superimposed on the head. Neck straightening in the apo state is incompatible with binding of MukE. This results in release of MukB from DNA-bound MukEF upon ATP hydrolysis, as illustrated on the right. MukB remains attached via the cWHD and linker of MukF. (**B**) Comparison of the inferred post-loading holding state of MukBEF and the post-extrusion holding state of *E. coli* Wadjet I (PDB: 8Q72). Models were superimposed on the KITE subunits MukE/JetB of the colored monomer. The second monomer is shown in transparent gray. The captured DNA in the post-loading state corresponds to the extruded loop in the post-extrusion state. (**C**) Model of the MukBEF activity cycle as in **Figure 5E**. Experimental structures and tentative models are shown in the bottom row. The tentative loop insertion state is shown in transparent colors, with experimental sub-structures highlighted in full color. Three-dimensional models for the tentative states are available in **Data S4**. See also **Figure 5** and **Data S4**.

## METHODS

### RESOURCE AVAILABILITY

#### Lead contact

Further information and requests for resources and reagents should be directed to and will be fulfilled by the lead contact, Jan Löwe.

#### Materials availability

All unique reagents generated in this study are available upon request, restricted by the use of a material transfer agreement (MTA).

#### Data and code availability

EM density maps have been deposited in the EMDB. Atom coordinates have been deposited in the PDB. Proteomics data, raw gel images and light microscopy images with associated analysis files have been deposited at Zenodo. The deposited data will be available as of the date of publication. Accession numbers will be listed in the key resources table. All other data will be available upon request. Any additional information required to reanalyze the data reported in this paper will be available from the lead contact upon request.

### EXPERIMENTAL SUBJECT AND MODEL DETAILS

#### *E. coli* strains

Strains are based on *E. coli* MG1655 and are listed in **Table S1**. The parental strain was obtained from the DSMZ strain collection (DSM 18039). All strains were viable in LB media at 37 °C, except for Δ*muk* strains and strains expressing *mukB(E1407Q)*, which were grown at 22 °C. Strains were single-colony purified and verified by marker analysis, PCR, and Sanger sequencing as required. Pre-cultures for all experiments were grown side-by-side to stationary phase and used freshly. Proteins were purified from *E. coli* BL21(DE), BL21-Gold(DE3), or *E. coli* C41(DE3), transformed with the appropriate expression plasmids as indicated (see also **Table S2** and **Data S5**).

### METHOD DETAILS

#### Genome engineering for strain construction

Replacement of the endogenous *mukFEB* locus in *E. coli* by its *P. thracensis* version was performed using a CONEXER-based strategy as described ^18,70^. Briefly, the *P. thracensis mukFEB* locus containing a HaloTag on *mukB* and a kanamycin resistance cassette was assembled into pFB411 containing *oriT* and a crDNA locus targeting the sites flanking the insert. The assembly reaction was transformed into donor strain SFB065 carrying the mobilizer plasmid pJF146. The acceptor strain SFB053 Δ*mukFEB::pheS(T251A, A294G) hygR* carrying the recombination plasmid pKW20 with *cas9* and *λ-red* under an arabinose-inducible promoter was induced in LB media with 5 μg/mL tetracycline and 0.5 % L-arabinose for 1 h at 37 °C. Donor and acceptor were mixed, and conjugation was performed for 1 h on TYE agar at 30 °C. Recombination was performed in LB media with 12.5 μg/mL kanamycin for 1 h at 37 °C followed by 18 h at 22 °C. Cultures were then plated on LB with 2 % glucose, 12.5 μg/mL kanamycin and 2.5 mM 4-chloro phenylalanine. Plates were incubated at 22 °C until colonies appeared. The annotated sequence of the modified locus is available in **Data S5**.

#### Protein production and purification

All protein concentrations were determined by absorbance at 280 nm using theoretical absorption coefficients. Annotated sequences of expression constructs are provided in **Data S5**. See also **Table S2**.

##### GST-hSENP1

GST-tagged hSENP1 protease was produced from a T7 expression plasmid (pFB83) in *E. coli* C41(DE3) by induction with 1 mM IPTG in 2xYT medium at 18 °C overnight. All purification steps were carried out at 4 °C. 83 g of cells were resuspended in 300 mL of buffer A (50 mM Tris/HCl pH 8.0 at room temperature (RT), 150 mM NaCl, 1 mM EDTA pH 8 at RT, 5 % glycerol, 2 mM DTT) supplemented with protease inhibitor cocktail (Roche) and Benzonase (Merck) and lysed at 172 MPa in a high-pressure homogenizer. The lysate was cleared by centrifugation at 40,000 x g for 30 min and incubated with 10 mL Glutathione Sepharose 4B (GE Healthcare) for 14 h. The resin washed with 15 column volumes (CV) of buffer A, 5 CV of buffer B (50 mM Tris/HCl pH 8.0 at RT, 500 mM NaCl, 1 mM EDTA pH 8 at RT, 5 % glycerol, 2 mM DTT) and protein was eluted in 5 CV of buffer A containing 3 mg/mL glutathione. Aliquots of the eluate were passed through a 0.22 µm filter and injected into a HiPrep 26/60 Sephacryl S-200 column (GE Healthcare) in buffer G1 (25 mM Tris/HCl pH 8 at RT, 250 mM NaCl, 0.5 mM DTT). Peak fractions were pooled, concentrated to 9.3 mg/mL on a Vivaspin 20 MWCO 30 filter (Sartorius), aliquoted, frozen in liquid nitrogen and stored at −80 °C.

##### MukBEF for loading assays and structural studies

*P. thracensis* MukBEF (NCBI accession identifiers WP_046975681.1, WP_046975682.1, and WP_046975683.1) was produced as described previously^18^ from a polycistronic expression construct assembled into a pET28 based backbone by Golden Gate cloning ^71^ (plasmids used: WT, pFB403; Cysteine mutant, pFB520 with MukF(D227C, Q412C) and MukB(R143C, R771C, C1118S, K1246C)). The construct contained a His_6_-SUMO tag fused to residue 1 of MukB which allowed affinity purification and scar-less tag removal by hSENP1 protease ^72^. The complex was produced in *E. coli* BL21-Gold(DE3) by autoinduction in ZYP-5052 media ^73^ at 24 °C. All purification steps were carried out at 4 °C. 15 g of cells were resuspended in 90 mL of IMAC buffer (50 mM Tris, 300 mM NaCl, 40 mM imidazole, 1 mM TCEP, pH 7.4 at RT) supplemented with protease inhibitor cocktail and Benzonase and lysed at 172 MPa in a high-pressure homogenizer. The lysate was cleared by centrifugation at 96,000 x g for 30 min, passed through a 0.45 μm filter, and incubated for 30 min with 25 mL Ni-NTA agarose (Qiagen) equilibrated in IMAC buffer. The resin was packed into a gravity flow column and washed with 3 x 50 mL IMAC buffer, then resuspended in 25 mL IMAC buffer containing 1 mg GST-hSENP1 and incubated for 1 h on a roller. The eluate was collected and pooled with a 12.5 mL wash using IMAC buffer, diluted with 18.8 mL buffer Q (10 mM Tris, pH 7.4 at RT), passed through a 0.22 μm filter and applied to a 20 mL HiTrap Heparin HP column (GE Healthcare). MukBEF was largely found in the flowthrough and was applied to a 5 mL HiTrap Q HP column (GE Healthcare). The column was washed with 2 CV of 10 mM Tris, 200 mM NaCl, 1 mM TCEP, pH 7.4 at RT, and protein was eluted with a 20 CV linear gradient from 200 mM NaCl to 1 M NaCl in buffer Q. MukBEF eluted at about 450 mM NaCl, was concentrated to 0.5 mL on a Vivaspin 20 MWCO 30 filter and was injected into a Superose 6 Increase 10/300 GL column (GE Healthcare) in buffer H200 (20 mM Hepes, 200 mM NaCl, 1 mM TCEP, pH 7.3 at RT). Peak fractions were pooled, concentrated to 6-9 mg/mL on a Vivaspin 2 MWCO 30 filter, aliquoted, frozen in liquid nitrogen and stored at −80 °C until use.

*P. thracensis* MukB was produced from pFB468 and purified as above except for omission of the Heparin step.

Due to its toxicity, cysteine mutant *P. thracensis* MukBEF^EQ^ was reconstituted in extracts by co-lysis of cells producing MukBC^Cys, EQ^ (pFB525) and MukEF^Cys^ (pFB522), respectively, as described ^18^. The His_6_-SUMO-MukB^EQ^ construct was propagated and produced at 22 °C. Cell pellets of both strains (15 g each) were mixed in 180 mL IMAC buffer, and the complex was purified as the wild-type construct.

Cysteine mutant *P. thracensis* MukB^EQ^ was purified as above except for omission of the Heparin step.

Cysteine mutant *E. coli* MukBEF (NCBI accession identifiers NP_415442.1, NP_415443.2, and NP_415444.1) and MukB were produced from pFB661 and pFB662, respectively, and were purified exactly as *P. thracensis* MukBEF, including the heparin step. The mutant complex contained MukB(R143C, R771C, C1118S, K1246C) and MukF(D227C, Q412C).

##### MukEF for SEC and structural studies

*E. coli* MukEF was produced from a bicistronic vector (pFB69) with a His_6_-SUMO tag fused to residue 1 of MukE. The complex was produced in *E. coli* BL21-Gold(DE3) by autoinduction in ZYP-5052 media ^73^ at 24 °C. All purification steps were carried out at 4 °C. 35 g of cells were resuspended in 175 mL of IMAC buffer (50 mM Tris, 300 mM NaCl, 20 mM imidazole, 1 mM TCEP, pH 7.4 at RT) supplemented with protease inhibitor cocktail and Benzonase and lysed at 172 MPa in a high-pressure homogenizer. The lysate was cleared by centrifugation at 96,000 x g for 30 min, passed through a 0.45 μm filter, and incubated for 30 min with 25 mL Ni-NTA agarose (Qiagen) equilibrated in IMAC buffer. The resin was packed into a gravity flow column and washed with 3 x 50 mL IMAC buffer, then resuspended in 25 mL SENP buffer (10 mM sodium phosphate, 50 mM NaCl, 20 mM imidazole, pH 7.4 at RT) containing 1 mg GST-hSENP1 and incubated for 1:45 h on a roller. The eluate was collected and pooled with a 12.5 mL wash using IMAC buffer, and 35 mL were mixed with 100 mL buffer Q (10 mM Tris, 50 mM NaCl, pH 7.4 at RT), passed through a 0.22 μm filter and applied to a 5 mL HiTrap Q HP column (GE Healthcare). The column was washed with 2 CV of buffer Q, and protein was eluted with a 20 CV linear gradient to 1 M NaCl in buffer Q. MukEF eluted at about 450 mM NaCl. Peak fractions were pooled and injected into a Sephacryl S-200 26/60 column in SEC buffer (10 mM Tris, 200 mM NaCl, 1 mM TCEP, 1 mM NaN_3_). Peak fractions were pooled and concentrated in a Vivaspin 20 MWCO 10 filter to 12 mg/mL, aliquoted, frozen in liquid nitrogen and stored at −80 °C until use.

*P. thracensis* MukEF (pFB481) was purified in an identical way, with the following exceptions: 60 g of cells were resuspended in 250 mL IMAC buffer containing 40 mM imidazole, hSENP1 digestion was done in IMAC buffer, and the SEC buffer was 20 mM HEPES, 200 mM NaCl, 1 mM TCEP, pH 7.3 at RT.

##### MukBEF subunits for pull-down assays

Hexa-histidine tagged *E. coli* MukB was overexpressed using the T7/pET system in BL21(DE3) cells using a pET21-MukB^his^ vector (gift from Gemma Fisher, MRC LMS). Cells were grown in LB supplemented with ampicillin to an OD_600_ value of 0.5-0.6, then induced with 1 mM IPTG and grown for a further 3 hours. Cells were then harvested by centrifugation and resuspended in 50 mM Tris-Cl pH 7.5, 250 mM NaCl, 1 mM DTT, 1 mM EDTA, 10 % sucrose. The cells were sonicated following addition of 0.01 mg/mL DNase I and 1 mM MgCl_2_ and the cell extract obtained by centrifugation. MukB was purified using a HisTrap affinity column (Cytiva). The column was equilibrated in buffer A (20 mM HEPES-KOH pH 7.7, 300 mM NaCl, 20 mM imidazole) and eluted with a 10 CV gradient from 50 to 400 mM imidazole. Peak fractions were pooled and dialyzed overnight against buffer C (20 mM HEPES-KOH pH 7.7, 50 mM NaCl, 2 mM EDTA, 1 mM DTT, 5% glycerol). MukB was further purified using a HiTrap Heparin column. The column was equilibrated in buffer C and eluted with a 16 CV gradient from 50 to 800 mM NaCl. MukB eluted in two peaks and the ‘low salt’ and ‘high salt’ samples were pooled separately. The ‘high salt’ sample was dialyzed against 20 mM HEPES–KOH pH 7.7, 200 mM NaCl, 5% glycerol, 1 mM EDTA, 1 mM DTT. Protein concentration was determined using a theoretical extinction co-efficient. The protein was frozen in liquid nitrogen and stored at −80 °C.

Purified MukF protein was a gift from Gemma Fisher (MRC LMS). MukE and MukEF complex were overexpressed as CPD fusion proteins using pFB062 and pFB070 respectively (see **Table S2**) transformed into BL21(DE3) cells. Transformed cells were grown at 37 °C in LB supplemented with kanamycin to an OD_600_ value of 0.4. MukE or MukEF expression was then induced by addition of 0.4 mM IPTG for 3 h at 25 °C. Cells were harvested by centrifugation, resuspended in lysis buffer (0.5 mM NaCl, 50 mM Tris-Cl pH 7.5, 15 mM imidazole, 10% glycerol) and flash frozen in liquid nitrogen. Cell suspensions were thawed, lysed by sonication and cleared by centrifugation. The His-tagged CPD fusion proteins were then purified as follows. Ni-NTA Agarose beads (Qiagen) were equilibrated with lysis buffer, before the fusion proteins were added and incubated for 2 h at 4 °C with gently shaking. The agarose beads were pelleted at 2,000 g and the supernatant removed. The pellet was washed three times with lysis buffer to remove unbound proteins. The self-cleavage activity of CPD was induced by the addition of 50 µM inositol hexakisphosphate (Sigma Aldrich), and the cleavage reaction allowed to proceed at 25 °C for 2 h with gentle shaking. Beads were pelleted and supernatant containing cleaved MukE or MukEF was removed. Protein was dialyzed against 50 mM Tris-HCl, pH 7.5, 50 mM NaCl, 0.1 mM DTT, 0.1 EDTA overnight and then further purified by ion exchange chromatography using a MonoQ column (Cytiva) equilibrated in the dialysis buffer. Protein was eluted by applying a salt gradient from 50 – 1000 mM NaCl over 30 CV. Peak fractions were pooled and dialyzed overnight against 50 mM Tris-HCl pH 7.5, 200 mM NaCl, 0.1 mM DTT, 0.1 M EDTA, 10 % glycerol. Concentrations were determined using theoretical extinction coefficients and proteins were stored at −80 °C.

##### Topoisomerase IV

*P. thracensis* ParE and ParC (NCBI accession identifiers AKH64223.1 and AKH64224.1) were cloned separately as His_6_-SUMO fusions into a pET28 based backbone by Golden Gate cloning (pFB478, ParE; pFB479, ParC). Proteins were produced in *E. coli* BL21-Gold(DE3) by autoinduction in ZYP-5052 media ^73^ at 24 °C. All purification steps were carried out at 4 °C and were identical for both proteins. 15 g of cells were resuspended in 90 mL of IMAC buffer (20 mM HEPES/KOH, 800 mM NaCl, 40 mM Imidazole, 1 mM TCEP, 10 % glycerol, pH 7.5 at RT) supplemented with protease inhibitor cocktail and lysed at 172 MPa in a high-pressure homogenizer. The lysate was then cleared by centrifugation at 96,000 x g for 30 min, sonicated to reduce viscosity, passed through a 0.45 μm filter, and incubated for 30 min with 2.5 mL Ni-NTA agarose (Qiagen) equilibrated in IMAC buffer. The resin was packed into a gravity flow column and washed with 2 x 25 mL IMAC buffer, 1x 25 mL SENP buffer (20 mM HEPES/KOH, 300 mM NaCl, 40 mM Imidazole, 1 mM TCEP, 10 % Glycerol, pH 7.5 at RT), then resuspended in 15 mL SENP buffer containing 1 mg GST-hSENP1 and incubated for 1 h on a roller. The eluate was passed through a 0.22 μm filter and applied to Sephacryl S-200 26/60 column (GE Healthcare) in SEC buffer (20 mM HEPES/KOH, 200 mM NaCl, 1 mM TCEP, 10 % glycerol, pH 7.5 at RT). Peak fractions were pooled, concentrated to 13-17 mg/mL on a Vivaspin 20 MWCO 30 filter, aliquoted, frozen in liquid nitrogen and stored at −80 °C until use. The Topo IV holoenzyme was reconstituted at 50 μM in SEC buffer by incubating an equimolar mixture of ParE and ParC for 1 h on ice. The reconstituted enzyme was then aliquoted, frozen in liquid nitrogen and stored at −80 °C until use.

##### DNA gyrase

*E. coli* GyrA and GyrB (NCBI accession identifiers NP_416734.1 and YP_026241.1) were cloned separately as His_6_-SUMO fusions into a pET28 based backbone by Golden Gate cloning (pFB638, GyrA; pFB639, GyrB). Proteins were produced in *E. coli* BL21-Gold(DE3) by autoinduction in ZYP-5052 media ^73^ at 24 °C. All purification steps were carried out at 4 °C and were identical to the purification of the Topo IV subunits, with the following modifications. After SEC, peak fractions were pooled and applied to a 1 mL HiTrap Q HP (GE Healthcare) in SEC buffer, washed with SEC buffer, and eluted with a 20 CV gradient into 50 % QB buffer (20 mM HEPES/KOH, 1 M NaCl, 1 mM TCEP, 10 % glycerol, pH 7.5 at RT). Peak fractions were pooled, concentrated to 10-20 mg/mL on a Vivaspin 2 MWCO 10 filter, aliquoted, frozen in liquid nitrogen and stored at −80 °C until use. The gyrase holoenzyme was reconstituted at 25 μM in SEC buffer by incubating an equimolar mixture of GyrA and GyrB on ice for 1 h. The reconstituted enzyme was then aliquoted, frozen in liquid nitrogen and stored at −80 °C until use.

##### gp5.9

T7 gp5.9 was produced from insect cells with modifications to a method described previously^42^. Briefly, Hi5 cells were infected with P3 virus and incubated for 72 h at 27 °C with shaking before cells were harvested by centrifugation. The pellet from a 2 L culture was resuspended in 100 mL lysis buffer (20 mM Tris-HCl pH 7.5, 200 mM NaCl, 2 mM β-mercaptoethanol, 10 % glycerol, protease inhibitor cocktail (Roche, as directed by the manufacturer), 20 mM imidazole). The cells were lysed by sonication and centrifuged to remove cell debris. The supernatant was then applied to Talon resin (Takara Bio) to purify gp5.9 using the histidine tag. Beads were equilibrated by washing three times with 15 mL wash buffer (20 mM Tris-HCl pH 7.5, 200 mM NaCl, 2 mM β-mercaptoethanol, 10 % glycerol, 20 mM imidazole). Supernatant from the centrifuged cell lysate was added to the beads and incubated for 30 min at 4 °C. The beads were then spun down and the supernatant (unbound protein) was removed. The beads were washed four times with wash buffer before gp5.9 was eluted with 50 mL elution buffer (20 mM Tris-HCl pH 7.5, 200 mM NaCl, 2 mM β-mercaptoethanol, 200 mM imidazole). The protein was next cleaved by adding 3C protease (Takara Bio, concentration as directed by the manufacturer) and incubating for 30 min, followed by dialysis against 20 mM Tris-HCl pH 7.5, 200 mM NaCl, 2 mM β-mercaptoethanol to remove imidazole. The sample was next passed over a 5 mL HisTrap HP column (Cytiva) to remove the cleaved tag and uncleaved gp5.9. The free gp5.9 in the flowthrough was loaded onto a 1 mL MonoQ column (Cytiva) equilibrated in buffer A (20 mM Tris-HCl pH 7.5, 1 mM TCEP, 100 mM NaCl) and was eluted with a gradient to buffer B (20 mM Tris-HCl pH 7.5, 1 mM TCEP, 1 M NaCl). Peak fractions were pooled and dialyzed against 20 mM Tris-HCl pH 7.5, 1 mM TCEP, 200 mM NaCl. The concentration of gp5.9 was calculated using a theoretical extinction coefficient of 8480 M^-1^ cm^-1^. The final protein was flash frozen and stored at –80 °C following supplementation with glycerol to 10 % final concentration.

#### DNA substrates

Plasmid substrates were pUC19 (2686 bp) or pFB526/pFB527 (both 2124 bp), which are a shortened versions of pUC19 lacking the *lacZ*α region. Negatively supercoiled DNA was prepared from overnight cultures of DH5α grown in LB media with 100 μg/mL ampicillin at 37 °C, and was purified using a QIAprep Spin miniprep or HiSpeed Plasmid Maxi kit (Qiagen). DNA was nicked with Nb.*Bts*I (NEB) or relaxed with *E. coli* Topo I (NEB) as recommended by the manufacturer and purified using a QIAquick PCR Purification kit (Qiagen).

#### BMOE cross-linking

Cysteine mutant *P. thracensis* MukBEF dimers were mixed at 1 μM with 6 ng/μL of negatively supercoiled pFB527 in SEC buffer (20 mM HEPES, 200 mM NaCl, 1 mM TCEP, pH 7.3 at RT) and incubated for 5 min on ice. The sample was then mixed with an equal amount of dilution buffer (20 mM HEPES, 30 mM NaCl, 1 mM TCEP, pH 7.3 at RT) and passed through a Zeba spin column (Thermo Fisher) in dilution buffer containing 1 mM ATP (pH 7.4), 2 mM MgCl_2_ and 0.05 % β-octyl glucoside. The sample was incubated at 22 °C for 1 h, after which 0.5 mM BMOE was added. The sample was incubated for 1 min, mixed with LDS-PAGE loading dye (Thermo Fisher) at a final concentration of 1 % 2-mercaptoethanol, incubated at 95 °C for 5 min, and resolved on a 4-16% Bis-Tris NuPAGE gel (Thermo Fisher) before Coomassie staining.

#### DNA entrapment assays

Cysteine mutant MukBEF dimers were mixed at 150 nM with 6 ng/μL plasmid DNA in loading buffer (10 mM Bis-Tris-Propane/HCl, 10 mM MgCl_2_, 0.1 mM TCEP, pH 7.0) containing 5 mM ATP/pH 7.4, or an ATP regeneration system (1 mM ATP/pH 7.4, 3 mM phosphoenolpyruvate, 1 mM NADH, 13 U/mL pyruvate kinase/lactate dehydrogenase) where indicated. Under standard low-salt conditions the reactions contained less than 5 mM NaCl carried over from the protein preparations. Reactions were incubated for the indicated times at 22 °C, and then cross-linked with 0.5 mM BMOE for 1 min. Where indicated, samples were treated with 0.2 U/μL of Nb.*Bts*I for 10 min at 37 °C after cross-linking to make their electrophoretic mobility comparable. Samples were mixed with Purple Gel Loading Dye (NEB) at a final concentration of 0.08 % SDS and resolved on 0.8 % agarose gels in 0.5x TBE buffer. Gels contained SYBR Safe DNA Gel Stain (Thermo Fisher) at 10,000x dilution suggested by the manufacturer.

Entrapment assays in the presence of topoisomerases were performed as indicated above but contained a final concentration of 30 mM NaCl. Topoisomerases were buffer exchanged into SEC buffer (20 mM HEPES, 200 mM NaCl, 1 mM TCEP, pH 7.3 at RT) immediately before use, and pre-mixed with MukBEF before dilution into the reaction mix. The final enzyme concentrations used were 100 nM Topo I, 50 nM Topo IV, and 50 nM GyrAB.

For inhibition assays with gp5.9, MukBEF was pre-mixed with gp5.9 at the indicated molar ratios and compensating volumes of gp5.9 buffer (20 mM Tris, 200 mM NaCl, 0.5 mM TCEP, 10 % glycerol, pH 7.4), or gp5.9 was added at the indicated timepoints after reaction start. Reactions were performed using nicked substrate and contained a final concentration of 12 mM NaCl carried over from the protein preparations.

#### Size-exclusion chromatography of gp5.9/MukEF

gp5.9 dimers at 15 μM final concentration were mixed on ice with MukE_4_F_2_ at 30 μM final concentration in SEC buffer (20 mM Tris, 200 mM NaCl, 0.5 mM TCEP, pH 7.4 at 22 °C) and injected into a Superose 6 Increase 3.2/300 column in SEC buffer. Chromatography was performed at 4 °C at a flow rate of 0.04 mL/min.

#### Cryo-EM sample preparation

##### DNA capture state

Wild-type *P. thracensis* MukBEF dimers at 150 nM were mixed in a total volume of 500 μL with 6 ng/μL nicked pFB526 in loading buffer (10 mM Bis-Tris-Propane/HCl pH 7.0, 10 mM MgCl_2_, 5 mM ATP/NaOH pH 7.4) and incubated for 1 h at RT. Optionally, 52.6 μL of 10 mM Na_3_VO_4_ in 50 mM Bis-Tris-Propane/HCl pH 7.0 or 26.3 μL of 10 mM BeSO_4_ / 200 mM NaF were added for a final concentration of 1 mM Na_3_VO_4_ or 0.5 mM BeSO_4_/10 mM NaF, respectively, and incubated for further 10 min at RT. The samples were then placed for 5 min on ice before concentration in a Vivaspin 500 30 k filter to 40-45 μL at 4 °C. The samples were kept on ice before application of 2.5 μL to UltrAuFoil m200 R2/2 grids that had been treated for 60 s at 35 mA in an Edwards glow discharger. The grids were immediately blotted using a Vitrobot Mark IV (FEI) operated at 4 °C and 100 % humidity and plunge-frozen in liquid ethane.

##### gp5.9/MukEF

An optimal ratio of gp5.9 to *E. coli* MukEF was found by SEC titration. For cryo-EM sample preparation, MukEF was mixed at 1 μM with 4 μM gp5.9 in buffer (20 mM Tris, 200 mM NaCl, 0.5 mM TCEP, 0.05 % b-octyl glucoside, pH 7.4 at 22 °C) and incubated on ice for 20 min. A volume of 2.5 μL was applied to a Quantifoil CuRh m200 R2/2 grid treated for 15 s at 30 mA in an Edwards glow discharger. The grid was immediately blotted using a Vitrobot Mark IV operated at 4 °C and 100 % humidity and plunge-frozen in liquid ethane.

#### Cryo-EM data collection

##### DNA capture state

Data was collected on three different grids in one continuous session: 1) ATP, 2) ATP/Na_3_VO_4_ and 3) ATP/BeF. Data was collected on a TFS Titan Krios with X-FEG emitter at 300 kV, equipped with a Gatan K3 detector operating in counting mode and a Gatan Quantum energy filter with 20 eV slit width centered on the zero-loss peak, and a 100 μm objective aperture inserted. Movies were acquired at four areas per hole using the aberration-free image shift (AFIS) method in EPU. The pixel size was 1.17 Å, the target defocus was −1 to −2.8 μm, and the total electron fluence was 40 e^-^/A^2^ collected over 2.8 s and fractionated into 40 frames.

##### gp5.9/MukEF

Data was collected on a single grid on a TFS Titan Krios with X-FEG emitter at 300 kV, equipped with a Gatan K3 detector operating in counting mode and a Gatan Quantum energy filter with 20 eV slit width centered on the zero-loss peak, and a 100 μm objective aperture inserted. Movies were acquired at four areas per hole using AFIS method in EPU. The pixel size was 0.928 Å, the target defocus was −1 to −2.4 μm, and the total electron fluence was 40 e^-^/A^2^ collected over 1.4 s and fractionated into 40 frames.

#### gp5.9 bacterial expression plasmids and toxicity tests

We have previously described the expression and purification of gp5.9 from insect cells and reported that gp5.9 toxicity prevented cloning and expression in *E. coli* using the T7/pET system ^42^. However, we found that we were able to maintain gp5.9 expression plasmids in *E. coli* using modified pBAD vectors containing the *rop* gene for very low copy number control and the tight induction control provided by the arabinose-inducible araBAD promoter ^74^. The gene encoding bacteriophage T7 gp5.9 (UniProt P20406) was ordered as a synthetic construct (GeneArt, Invitrogen) either without a tag or with a C-terminal FLAG tag flanked by *Eco*RI and *Hind*III restriction sites. These were cloned into the pBAD322K vector using standard techniques to form vectors expressing variants of gp5.9 named pBAD322K-gp5.9 and pBAD322K-gp5.9^FLAG^. The integrity of these constructs was confirmed by sequencing. To test for toxicity of gp5.9 expression the expression plasmids (25 ng each) were transformed into chemically-competent MG1655 or MEK1326 (Δ*recB*) cells before plating on agar containing LB + 50 μg/mL kanamycin, either with or without 1 % L-arabinose to induce expression of gp5.9.

For spot dilution tests of *mukFEB* modified strains, similar constructs with an ampicillin resistance cassette were used (pBAD322A and pBAD322A-gp5.9). Transformed strains were grown overnight in LB + 100 μg/mL ampicillin, diluted in LB, and then 7.5 μL of the dilutions were spotted on LB agar containing 100 μg/mL ampicillin and 1 % L-arabinose. Plates were incubated at 37 °C for 16 h.

#### gp5.9 pulldown proteomics

MG1655 and MEK1326 (Δ*recB*) *E. coli* were transformed with 50 ng of either pBADK-gp5.9 (for the mock condition) or pBADK-gp5.9^FLAG^ (for the pulldown condition), plated on LB agar plates containing 50 μg/mL kanamycin, and incubated overnight at 37 °C. LB/kanamycin overnight starter cultures were made for each condition and 2mL each was added to 1 L LB containing 50 μg/mL kanamycin with shaking at 37 °C. At OD_600_ between 0.3-0.4, 0.2 % arabinose was added to induce expression of gp5.9 or g5.9^FLAG^. 10 mL aliquots were taken at 2 h post-induction, placed on ice and then spun at 4000 rcf and 4 °C to pellet the cells. Supernatants were discarded and cells were resuspended in 200 μL of resuspension buffer (50 mM Tris-Cl pH 8, 200 mM NaCl, 10 % sucrose, 1 mM DTT). Resuspended cells were stored at –20 °C. The cells were thawed and 0.1 % Triton X-100, followed by 0.1 mg/mL lysozyme, was added. Lysis mixtures were shaken at room temperature for 30 min before 0.01 mg/mL DNase I and 1 mM MgCl_2_ were added. Mixtures were shaken for a further 10 min and then spun in a microcentrifuge for 10 min at maximum speed to obtain the soluble cell extract. 10 μL resin of resuspended anti-FLAG M2 magnetic beads (Sigma-Aldrich) were extracted and used for pulldowns from the cell extracts performed following manufacturer’s instructions with minor modifications. Beads were washed and equilibrated with 150 μL base buffer (50 mM Tris-Cl pH 8, 200 mM NaCl, 1 mM DTT), before cell extract was incubated for 60 min at room temperature, with gentle mixing every 10 min. Beads were then washed three times with 200 μL base buffer, or until A_280_ of the wash liquid was below 0.05.

For proteomics analysis of the pull-down samples, 15 μL base buffer was used to cover the beads. These samples were then spun down, placed on ice and delivered to the University of Bristol Proteomics Facility for analysis. Samples were subjected to tryptic digest and TMT tagging before nano-LC MS/MS was performed, followed by a Sequest search against the Uniprot *E. coli* K12 database supplemented with the pBAD322K open reading frames (including gp5.9) and a common contaminants database. Data was filtered using a 5 % false discovery rate cut-off and a maximum fold change of 1000. Data for the four conditions were compared as abundance ratios for two repeats each of MG1655 pulldown/mock and Δ*recB* pulldown/mock (where mock refers to a pulldown experiment performed with untagged gp5.9). Pooled data refers to a comparison of four repeats for pulldown/mock where the MG1655 and Δ*recB* data were combined. The significance (p value) of the difference between pulldown and mock experiments was determined by multiple non-parametric t-tests and the data were not corrected for multiple comparisons. Volcano plots were created by plotting log_2_ of the abundance ratio against log_10_ of the significance (p) of this change. The top hits for gp5.9 pulldown were ranked using Manhattan scores calculated in VolcaNoseR ^75^.

#### Light microscopy

Starter cultures of MG1655 or MEK1326 (Δ*recB*) containing pBAD322K vectors were prepared by inoculating 5 mL LB + 50 μg/mL kanamycin + 1 % glucose (to suppress expression of toxic gp5.9) and incubating overnight at 37 °C with shaking at 250 rpm. These overnight cultures were then diluted 500-fold into LB + 50 μg/mL kanamycin and incubated at 37 °C until an OD_600_ value of 0.2 was reached. Expression was then induced with 0.2 % L-arabinose (or H_2_O as a no arabinose control). Cells were grown for a further 3 h at either 37 °C or 22 °C before 1 mL aliquots were removed to ice for 30 min. The cells were spun at 15000 rpm for 2 min, resuspended in 0.5 mL PBS, spun again and resuspended in 0.5 mL PBS and 2 % paraformaldehyde. After a 30 min incubation at room temperature with occasional mixing, the cells were spun and resuspended in 0.5 mL PBS and 1 μg/mL DAPI. 5 μL of cell culture, followed by 20 μL of Vectashield Antifade Mounting Medium with DAPI (Vector Laboratories), was applied to a coverslip that was then inverted onto a slide. Cellular morphology and nucleoids were imaged by combined phase contrast and fluorescence using a widefield microscope at 40x magnification.

#### *In vitro* pull down of MukBEF subunits

MG1655 cells containing either pBAD322K or pBAD322K-gp5.9^FLAG^ were grown as described for the microscopy experiment but were induced at 0.D_600_ ∼0.5-0.6 and then incubated for 3 h at 37°C at 250 rpm. Cells were pelleted by centrifugation at 3000 g for 10 min and resuspended in 1 mL resuspension buffer (50 mM tris pH 8, 200 mM NaCl, 1 mM DTT and 10 % sucrose) per 100 mL culture. 1 mL of resuspended cells were mixed with 0.1% Triton X-100 and 0.2 mg/mL lysozyme and shaken for 30 min at room temperature. Addition of 0.01 mg/mL DNaseI and 1 mM MgCl_2_ was then followed by shaking at room temp for 10 min and centrifugation at 15000 rpm for 10 min. The supernatant (soluble fraction) was used to bait magnetic beads. 30 μL Pierce Anti-DYKDDDDK Magnetic Agarose (ThermoFisher Scientific) bead slurry was applied to a DynaMag™-2 Magnet (Invitrogen) and the supernatant was removed. The beads were washed twice with 200 µL P buffer (50 mM Tris-Cl pH 8, 150 mM NaCl, 1 mM EDTA, 1 mM DTT) followed by supernatant removal. 500 μL of the soluble cell extract was applied to the beads and incubated for 10 min before magnetization, supernatant removal and three P buffer washes. Each magnet application was for 1 min and beads were rotating at room temperature for all incubations. For interaction analyses, the gp5.9-baited beads were mixed with purified MukBEF prey proteins, pre-incubated in various combinations (200 µL containing 1 µM each of MukB_2_E_2_F, MukE_2_F, MukB_2_, MukE_2_ or MukF as indicated) for 10 min then washed twice as above. FLAG-tagged gp5.9 and interacting partners were then eluted by 30 min incubation with 25 µg FLAG peptide in 50 μL of P buffer. Samples for each fraction were analyzed by SDS page. Band densities were quantified using ImageJ and normalized to the intensity of the eluted gp5.9 band.

### QUANTIFICATION AND STATISTICAL ANALYSIS

#### Phylogenetic analysis

Representative sequences for the Wadjet group were obtained by iterative profile searches with manual curation. We downloaded 254,733 bacterial and 2,809 archaeal genomes with at least scaffold-level assemblies from the NCBI, and clustered the protein sequences at 80 % identity using MMseqs2 linclust ^76^. We then created initial search profiles for MksB, MksF, MksE and MksG using sequences from ^24^ after clustering and MAFFT alignment ^77^. Profile searches were performed with MMseqs2 against the clustered database, using the parameters –s 7.5 –max-seqs 100000. We then identified candidate operons containing co-directional genes that produced consecutive hits with the MksB, MksF and MksE profiles, and an optional flanking hit with the MksG profile. Candidate operons were kept that had MksB proteins larger than 890 amino acids (AA) and contained Walker motives, MksF proteins between 400 and 1200 AA, and MksE proteins between 150 and 800 AA. Refined profiles were then built and used for sequence searches with HMMSearch ^78^. We performed six iterations of search, operon inference and profile refinement, and discarded operons that were less than two genes away from the end of a contig to ensure that only fully sequenced operons were retained. Finally, we used AlphaFold2 ^79^ to predict the structures of proteins encoded directly up- and downstream of operons lacking an MksG hit, and visually inspected them to verify the absence of the MksG nuclease. Wadjet operons with subunit assignments are listed in **Data S1**.

For the inference of a phylogenetic tree, we included sequences for Smc and Smc1–6 from ^80^ and added Loki- and Thorarchaeal SMC sequences from a MMseqs2 search. Two full-length alignments were constructed with MAFFT: 1) Smc and Smc1–6, and 2) MksB. Regions for the N- and C-terminal head and the hinge were extracted using structures of *B. subtilis* Smc and *E. coli* MukB as a guide, re-aligned separately, trimmed and catenated to generate a single composite alignment. Columns in the composite alignment containing more than 30 % gaps were removed. A phylogenetic tree was then inferred with IQ-Tree2 ^81^ using fast bootstrapping (-B 1000) and the model setting -m Q.pfam+F+I+I+R10, which had been automatically selected in exploratory runs. The tree was visualized with iTOL ^82^. The composite alignment and tree are available in **Data S2**.

#### Cryo-EM data analysis

Motion correction and dose weighting was performed in RELION ^83^ with one patch per micrograph and on-the-fly gain correction. The contrast transfer function (CTF) was fitted with CTFFIND4 ^84^. Automated particle picking was performed with crYOLO ^85^. All further processing was done in RELION and cryoSPARC ^86^. Maps were rendered in ChimeraX ^87^. Data collection and map statistics are shown in **Table 1**.

**Table 1.** Cryo-EM data collection and model statistics.

|  | Heads core | Open gate<br>(focused) | Open gate<br>(monomer) | DNA capture | DNA capture<br>(dimer) | gp5.9/MuKEF | gp5.9/MuKEF<br>(focused) |
| --- | --- | --- | --- | --- | --- | --- | --- |
|  | EMD-51442<br>PDB 9GM6 | EMD-51444<br>PDB 9GM8 | EMD-51443<br>PDB 9GM7 | EMD-51445<br>PDB 9GM9 | EMD-51446<br>PDB 9GMA | EMD-51447<br>PDB 9GMB | EMD-51448<br>PDB 9GMD |
| **Data collection and processing** |  |  |  |  |  |  |  |
| Magnification | 81,000 |  |  |  |  | 105,000 |  |
| Voltage (kV) | 300 |  |  |  |  | 300 |  |
| Electron fluence (e<sup>-</sup>/Å<sup>2</sup>) | 40 |  |  |  |  | 40 |  |
| Defocus range (μm) | -1 to -2.8 |  |  |  |  | -1 to -2.4 |  |
| Pixel size (Å) | 1.17 |  |  |  |  | 0.928 |  |
| Symmetry imposed | C1 |  |  |  |  | C1 |  |
| Initial particle images (no.) | 4,460,000 (Total)<br>1,200,000 (ATP/Na<sub>3</sub>VO<sub>4</sub>)<br>1,500,000 (ATP/BeF)<br>1,760,000 (ATP) |  |  |  |  | 3,500,000 |  |
| Final particle images (no.) | 210,276 | 34,436 | 34,436 | 7,508 | 3,754 | 57,528 | 57,528 |
| Map resolution (Å) | 3.5 | 3.9 | 4.3 | 7.8 | 9.1 | 4.2 | 4.0 |
| FSC threshold | 0.143 | 0.143 | 0.143 | 0.143 | 0.143 | 0.143 | 0.143 |
| **Model** |  |  |  |  |  |  |  |
| Initial model used<br>(PDB code) | 7NZ2 | 9GM6,<br>AlphaFold2 | 9GM8, 7NZ2 | 9GM8 | 9GM7, 9GM9 | AlphaFold2,<br>8B1R | AlphaFold2,<br>8B1R |
| Model resolution (Å) | 3.7 | 4.2 | 7.2 | 8.5 | 7.3 | — | — |
| FSC threshold | 0.5 | 0.5 | 0.5 | 0.5 | 0.5 | — | — |
| Map sharpening B factor (Å<sup>2</sup>) | -40 | -92 | — | -40 | — | -80 | -80 |
| **Model composition** |  |  |  |  |  |  |  |
| Non-hydrogen atoms | 23,924 | 28,582 | 34,093 | 29,658 | 66,565 | 9,003 | 4,995 |
| Protein residues | 2,956 | 3,541 | 4,218 | 3,439 | 7,856 | 1,111 | 611 |
| Nucleic acid residues | — | — | — | 93 | 146 | — | — |
| Ligands | PNS: 2<br>ATP: 2<br>Mg: 2 | PNS: 2<br>ATP: 2<br>Mg: 2 | PNS: 2<br>ATP: 2<br>Mg: 2 | PNS: 2<br>ATP: 2<br>Mg: 2 | PNS: 4<br>ATP: 4<br>Mg: 4 | — | — |
| **R.m.s. deviations** |  |  |  |  |  |  |  |
| Bond lengths (Å) | 0.005 | 0.012 | 0.013 | 0.017 | 0.017 | 0.009 | 0.004 |
| Bond angles (°) | 1.095 | 1.474 | 1.645 | 1.886 | 1.871 | 1.148 | 0.615 |
| **Validation** |  |  |  |  |  |  |  |
| MolProbity score | 1.50 | 1.42 | 1.32 | 1.46 | 1.36 | 1.67 | 1.73 |
| Clashscore | 5.87 | 4.64 | 3.57 | 5.06 | 3.97 | 9.18 | 9.48 |
| Poor rotamers (%) | 0.16 | 0.82 | 0 | 0.91 | 0.5 | 0.51 | 0 |
| **Ramachandran plot** |  |  |  |  |  |  |  |
| Favored (%) | 96.93 | 96.90 | 96.98 | 96.84 | 96.95 | 96.91 | 96.46 |
| Allowed (%) | 3.07 | 3.10 | 3.02 | 3.16 | 3.05 | 3.09 | 3.54 |
| Disallowed (%) | 0 | 0 | 0 | 0 | 0 | 0 | 0 |

##### Open-gate state and DNA capture state

Particles were picked using a crYOLO model trained on apo-MukBEF ^18^. We obtained 1.2 M particles from 9,063 micrographs for the ATP/Na_3_VO_4_ dataset, 1.5 M particles from 10,704 micrographs for the ATP/BeF dataset, and 1.7 M particles from 10,031 micrographs for the ATP dataset. Subsets of particles were selected by multiple rounds 2D classification, which were analyzed by 3D classification in RELION using a low-pass filtered map of apo-MukBEF as a reference. This revealed the presence of the open-gate state in all datasets. We then pooled the particles from all datasets and processed them further as follows.

We performed non-uniform refinement in cryoSPARC followed by one round of 3D classification without alignment in RELION, two rounds of focused classification without alignment using a mask around the heads to select 210,000 particles that reconstructed good density in the core of the head module. All datasets contributed to the density, and reconstructions split by dataset showed similar densities for the bound nucleotides, which were modeled as MgATP (**Figure S2F**). The map was improved by Bayesian polishing split by dataset, by per-particle defocus refinement, and by focused refinement with local pose search and Blush regularization. This resulted in the head core map at 3.5 Å resolution. To improve the density of the open gate, we performed focused classification without alignment using a mask that incorporated the gate. A subset of 34,000 particles was selected for refinement with local pose search and Blush regularization. This resulted in the open-gate map at 3.9 Å resolution. The MukBEF monomer was reconstructed from the same particles using flexible refinement in cryoSPARC. This resulted in the open-gate monomer map at 4.3 Å nominal resolution. The DNA capture state was obtained by further 3D classification in cryoSPARC using a threshold resolution of 9 Å. This selected 3,750 particles that reconstructed clear density for DNA. Re-centering on the DNA-bound gate and refinement revealed the dimeric nature of the capture state, yielding the dimer map at 9.1 Å nominal resolution. The map was then refined with C2 symmetry imposed, and the particle set was expanded in C2 to 7,500 particles. Particles were re-centered on the monomer, and the capture state was refined in C1 with local pose search and a mask around the head module and DNA binding site. This resulted in the DNA capture state map at 7.8 Å nominal resolution.

##### gp5.9/MukEF

Particles were automatically picked using a crYOLO model trained on manually picked examples. Subsets of particles were selected by two rounds of 2D classification and were then subjected to 3D classification in Relion using an initial model based on a MukEF crystal structure (PDB: 3EUH) filtered to 60 Å resolution. Selected particles were then subjected to *ab initio* reconstruction and 3D classification in cryoSPARC. This was followed by non-uniform refinement with C2 symmetry imposed, and symmetry expansion in C2. The structure was then refined without symmetry using a mask around one MukEF monomer, using local pose search with an alignment threshold of 6 Å. The gp5.9 protein was not visible at this stage but became apparent after one round of 3D classification in cryoSPARC. Particles were subsequently subjected to Bayesian polishing in Relion. We encountered occasional flipping of particles during local refinements, and thus reinstated the dataset to C1. Next, we refined the structure with global pose search using Blush regularization, yielding the gp5.9/MukEF map at 4.1 Å nominal resolution. We then subtracted the signal of the MukF core, and re-centered on the gp5.9/MukE region. This was subjected to a final focused refinement with global pose search and Blush regularization, yielding the gp5.9/MukEF focus map with improved gp5.9 density at 4 Å nominal resolution.

#### Structural model building

Map sharpening was performed by B-factor compensation and FSC weighting ^88^ where indicated. Starting models were obtained from the PDB or generated in AlphaFold2 ^79^, and model building and refinement was performed with ISOLDE ^89^, COOT ^90^ and phenix.real_space_refine ^91^. Model statistics were calculated with Phenix and are listed in **Table 1**.

##### Open-gate state and DNA capture state

The coiled-coil arms of PDB: 7NZ2 were flexibly fit into the open gate monomer map using ISOLDE, and then annealed into the head core map. The head module was built from fragments of 7NZ2 annealed into the head core map, whereby building of the ν-MukB larynx region was facilitated by an auxiliary map obtained by focused classification of this area. The model was then trimmed to the region of interest and subjected to a single macro-cycle in phenix.real_space_refine with restraints for the prosthetic group phosphopantetheine, secondary structure restraints, and Ramachandran restraints. Finally, AcpP, but not its prosthetic group, was replaced by chains G and h of PDB: 7NYW. This yielded the head core model.

To generate the open gate model, the head core model was rigid-body fit into the open gate map, together with an AlphaFold2 prediction of the MukF MD and nWHD regions. The model was adjusted by flexible fitting in ISOLDE.

The monomer model was generated by rigid-body fitting the open gate model into the monomer map and extending the coiled-coil arms with a model built into the monomer map as described above. The transition in the arm region was adjusted in ISOLDE.

The DNA capture state model was based on the open gate model and built into the capture state map. We generated a stretch of ideal B-form DNA in COOT using a sequence derived from the plasmid substrate. This was flexibly fit in ISOLDE using a κ value of 50. MukF was slightly adjusted, and the DNA interface was relaxed in ISOLDE using a κ value of 50. The dimeric capture state was obtained by extending the capture state model through rigid-body fitting into the capture state dimer map.

##### gp5.9/MukEF

A model for MukEF was generated in AlphaFold2. This was composed with gp5.9 in its RecBCD-bound form (PDB: 8B1R) by rigid-body fitting into the sharpened gp5.9/MukEF focus map, which had the best density for gp5.9. The composite model was then flexibly fitted in ISOLDE ^89^ with distance and torsion restraints, and local adjustments with relaxed restraints. Next, the model was refined in phenix.real_space_refine with secondary structure and Ramachandran restraints. In a parallel approach, the same strategy was applied to build into the sharpened non-focused gp5.9/MukEF map, which showed good density for the MukF MD and nWHD. We then merged the MD and nWHD from the non-focused model into the focused model, re-build the transition in ISOLDE, trimmed the model, and subjected it to phenix.real_space_refine with secondary structure and Ramachandran restraints to generate the final focused model. The final non-focused model was obtained by merging the final focused model into the working model, re-building the transition in ISOLDE, and subjecting the structure to refinement in phenix.real_space_refine with secondary structure and Ramachandran restraints.

#### Light microscopy image analysis

Images were analyzed using the FIJI Modular Image Analysis (MIA) plugin ^92^ with a custom workflow (DOI: 10.5281/zenodo.13748172). Detection of bacterial cells used a threshold of 0.5 μm length and erroneous cell selections were removed prior to statistical analysis.

## Supplemental Material

**Table S1.**
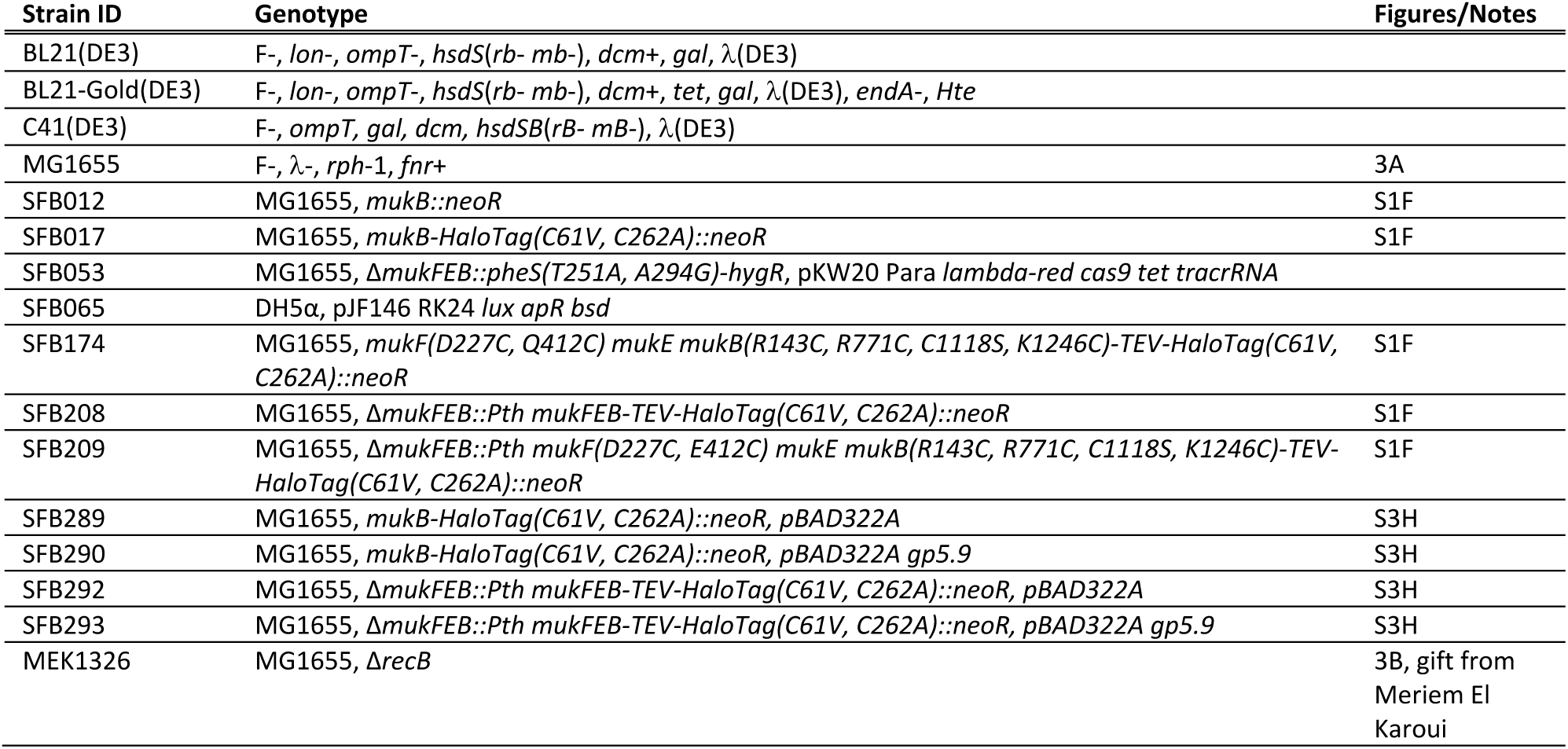
Bacterial strains.

**Table S2.** Plasmids.

| ID | Name | Description | Source |
| --- | --- | --- | --- |
| pFB062 | pET-Gate2 MukE-CPD-His10 | T7 expression plasmid for producing *E. coli* MukE | This study |
| pFB069 | pET-Gate2 MukF His6-SUMO-MukE | T7 expression plasmid for producing *E. coli* MukEF | This study |
| pFB070 | pET-Gate2 MukF MukE-CPD-His10 | T7 expression plasmid for producing *E. coli* MukEF | This study |
| pFB083 | pGEX GST-hSENP1 | T7 expression plasmid for producing GST-tagged hSENP1 | Komander lab |
| pFB403 | pET-Gate2 *Pth* MukF MukE His6-SUMO-MukB | T7 expression plasmid for producing SUMO-tagged *P. thracensis* MukBEF | Bürmann et al., 2021 |
| pFB411 | pCONEX-Gate4 CRISPR(mukFEB cloDF13) *ccdB* | Shuttle plasmid for targeting of the *mukFEB* locus (*BsaI* acceptor); crRNA targets pKW20 plasmid | Bürmann et al., 2021 |
| pFB468 | pET-Gate2 *Pth* His6-SUMO-MukB | T7 expression plasmid for producing *P. thracensis* MukB | Bürmann et al., 2021 |
| pFB478 | pET-Gate2 *Pth* His6-SUMO-ParE | T7 expression plasmid for producing *P. thracensis* ParE | This study |
| pFB479 | pET-Gate2 *Pth* His6-SUMO-ParC | T7 expression plasmid for producing *P. thracensis* ParC | This study |
| pFB481 | pET-Gate2 *Pth* MukF His6-SUMO-MukE | T7 expression plasmid for producing *P. thracensis* MukEF | This study |
| pFB520 | pET-Gate2 *Pth* MukF(D227C, Q412C) MukE His6-SUMO-MukB(R143C, R771C, C1118S, K1246C) | T7 expression plasmid for producing cysteine mutant *P. thracensis* MukBEF | This study |
| pFB522 | pET-Gate2 *Pth* MukF(D227C, Q412C) MukE | T7 expression plasmid for producing cysteine mutant *P. thracensis* MukEF | This study |
| pFB525 | pET-Gate2 *Pth* His6-SUMO-MukB(R143C, R771C, C1118S, K1246C, E1407Q) | T7 expression plasmid for producing cysteine mutant *P. thracensis* MukB(EQ) | This study |
| pFB526 | pUC19 matS2 | Entrapment assay substrate | This study |
| pFB527 | pUC19 matS2(scrambled) | Entrapment assay substrate | This study |
| pFB638 | pET-Gate2 His6-SUMO-GyrA | T7 expression plasmid for producing *E. coli* GyrA | This study |
| pFB639 | pET-Gate2 His6-SUMO-GyrB | T7 expression plasmid for producing *E. coli* GyrB | This study |
| pFB661 | pET-Gate2 MukF(D227C, Q412C) MukE His6-SUMO-MukB(R143C, R771C, C1118S, K1246C) | T7 expression plasmid for producing cysteine mutant *E. coli* MukBEF | This study |
| pFB662 | pET-Gate2 His6-SUMO-MukB(R143C, R771C, C1118S, K1246C) | T7 expression plasmid for producing cysteine mutant *E. coli* MukB | This study |
|  | pBAD322K | Arabinose inducible vector, *kanR* | Cronan, 2006 |
|  | pBAD322K-gp5.9 | Arabinose inducible gp5.9, *kanR* | This study |
|  | pBAD322K-gp5.9-FLAG | Arabinose inducible gp5.9-FLAG, *kanR* | This study |
|  | pBAD322A | Arabinose inducible vector, *ampR* | Cronan, 2006 |
|  | pBAD322A-gp5.9 | Arabinose inducible gp5.9, *ampR* | This study |
|  | pACEBac1 His6-3C-gp5.9 | Insect cell expression of gp5.9 | Wilkinson et al., 2022 |
|  | pET21a MukB-His6 | T7 expression plasmid for producing *E. coli* MukB | Zawadzka et al., 2018 |
| pUC19 |  | Entrapment assay substrate | New England Biolabs |
| pJF146 | RK24 *lux apR bsd* | RK2 conjugation machinery; NCBI: MK809154.1 | Fredens et al., 2019 |
| pKW20 | Para *lambda-red cas9 tet tracrRNA* | REXER helper plasmid; NCBI: MN927219.1 | Wang et al., 2016 |

## Supplemental Files

**Movie S1.** Model for DNA loading of MukBEF.

**Data S1**. Wadjet group operons.

**Data S2**. Composite alignment and phylogenetic tree of SMC proteins.

**Data S3**. TMT-MS analysis of gp5.9 pull-down experiments.

**Data S4**. Tentative models for holding and open ring states. The models were not built into experimental density and are intended to give a 3D impression of the pathway shown in **Figure 5E**.

**Data S5**. Annotated sequences of plasmids and genomic loci.

## References

1. Yatskevich, S., Rhodes, J., and Nasmyth, K. (2019). Organization of Chromosomal DNA by SMC Complexes. Annu. Rev. Genet. 53, 445–482. 10.1146/annurev-genet-112618-043633.

2. Bürmann, F., and Löwe, J. (2023). Structural biology of SMC complexes across the tree of life. Curr. Opin. Struct. Biol. 80, 102598. 10.1016/j.sbi.2023.102598.

3. Gruber, S. (2018). SMC complexes sweeping through the chromosome: going with the flow and against the tide. Curr. Opin. Microbiol. 42, 96–103. 10.1016/j.mib.2017.10.004.

4. Shintomi, K., Takahashi, T.S., and Hirano, T. (2015). Reconstitution of mitotic chromatids with a minimum set of purified factors. Nat Cell Biol 17, 1014–1023. 10.1038/ncb3187.

5. Houlard, M., Godwin, J., Metson, J., Lee, J., Hirano, T., and Nasmyth, K. (2015). Condensin confers the longitudinal rigidity of chromosomes. Nat Cell Biol 17, 771–781. 10.1038/ncb3167.

6. Ba, Z., Lou, J., Ye, A.Y., Dai, H.-Q., Dring, E.W., Lin, S.G., Jain, S., Kyritsis, N., Kieffer-Kwon, K.-R., Casellas, R., et al. (2020). CTCF orchestrates long-range cohesin-driven V(D)J recombinational scanning. Nature 586, 305–310. 10.1038/s41586-020-2578-0.

7. Le, T.B.K., Imakaev, M.V., Mirny, L.A., and Laub, M.T. (2013). High-resolution mapping of the spatial organization of a bacterial chromosome. Science 342, 731–734. 10.1126/science.1242059.

8. Lioy, V.S., Cournac, A., Marbouty, M., Duigou, S., Mozziconacci, J., Espéli, O., Boccard, F., and Koszul, R. (2018). Multiscale Structuring of the E. coli Chromosome by Nucleoid-Associated and Condensin Proteins. Cell 172, 771–783.e18. 10.1016/j.cell.2017.12.027.

9. Hoencamp, C., Dudchenko, O., Elbatsh, A.M.O., Brahmachari, S., Raaijmakers, J.A., van Schaik, T., Sedeño Cacciatore, Á., Contessoto, V.G., van Heesbeen, R.G.H.P., van den Broek, B., et al. (2021). 3D genomics across the tree of life reveals condensin II as a determinant of architecture type. Science 372, 984–989. 10.1126/science.abe2218.

10. Sjögren, C., and Nasmyth, K. (2001). Sister chromatid cohesion is required for postreplicative double-strand break repair in Saccharomyces cerevisiae. Curr Biol 11, 991–995.

11. Fujioka, Y., Kimata, Y., Nomaguchi, K., Watanabe, K., and Kohno, K. (2002). Identification of a novel non-structural maintenance of chromosomes (SMC) component of the SMC5-SMC6 complex involved in DNA repair. J Biol Chem 277, 21585–21591. 10.1074/jbc.M201523200.

12. Decorsière, A., Mueller, H., van Breugel, P.C., Abdul, F., Gerossier, L., Beran, R.K., Livingston, C.M., Niu, C., Fletcher, S.P., Hantz, O., et al. (2016). Hepatitis B virus X protein identifies the Smc5/6 complex as a host restriction factor. Nature 531, 386–389. 10.1038/nature17170.

13. Panas, M.W., Jain, P., Yang, H., Mitra, S., Biswas, D., Wattam, A.R., Letvin, N.L., and Jacobs, W.R., Jr (2014). Noncanonical SMC protein in Mycobacterium smegmatis restricts maintenance of Mycobacterium fortuitum plasmids. Proc Natl Acad Sci U A 111, 13264– 13271. 10.1073/pnas.1414207111.

14. Haering, C.H., Löwe, J., Hochwagen, A., and Nasmyth, K. (2002). Molecular architecture of SMC proteins and the yeast cohesin complex. Mol Cell 9, 773–788.

15. Haering, C.H., Farcas, A.-M., Arumugam, P., Metson, J., and Nasmyth, K. (2008). The cohesin ring concatenates sister DNA molecules. Nature 454, 297–301. 10.1038/nature07098.

16. Ivanov, D., and Nasmyth, K. (2005). A topological interaction between cohesin rings and a circular minichromosome. Cell 122, 849–860. 10.1016/j.cell.2005.07.018.

17. Wilhelm, L., Bürmann, F., Minnen, A., Shin, H.-C., Toseland, C.P., Oh, B.-H., and Gruber, S. (2015). SMC condensin entraps chromosomal DNA by an ATP hydrolysis dependent loading mechanism in Bacillus subtilis. Elife 4. 10.7554/eLife.06659.

18. Bürmann, F., Funke, L.F.H., Chin, J.W., and Löwe, J. (2021). Cryo-EM structure of MukBEF reveals DNA loop entrapment at chromosomal unloading sites. Mol. Cell 81, 4891–4906.e8. 10.1016/j.molcel.2021.10.011.

19. Shaltiel, I.A., Datta, S., Lecomte, L., Hassler, M., Kschonsak, M., Bravo, S., Stober, C., Ormanns, J., Eustermann, S., and Haering, C.H. (2022). A hold-and-feed mechanism drives directional DNA loop extrusion by condensin. Science 376, 1087–1094. 10.1126/science.abm4012.

20. Taschner, M., and Gruber, S. (2023). DNA segment capture by Smc5/6 holocomplexes. Nat. Struct. Mol. Biol. 30, 619–628. 10.1038/s41594-023-00956-2.

21. Niki, H., Jaffé, A., Imamura, R., Ogura, T., and Hiraga, S. (1991). The new gene mukB codes for a 177 kd protein with coiled-coil domains involved in chromosome partitioning of E. coli. EMBO J 10, 183–193.

22. Mäkelä, J., and Sherratt, D.J. (2020). Organization of the Escherichia coli Chromosome by a MukBEF Axial Core. Mol. Cell 78, 250–260.e5. 10.1016/j.molcel.2020.02.003.

23. Petrushenko, Z.M., She, W., and Rybenkov, V.V. (2011). A new family of bacterial condensins. Mol Microbiol 81, 881–896. 10.1111/j.1365-2958.2011.07763.x.

24. Doron, S., Melamed, S., Ofir, G., Leavitt, A., Lopatina, A., Keren, M., Amitai, G., and Sorek, R. (2018). Systematic discovery of antiphage defense systems in the microbial pangenome. Science 359. 10.1126/science.aar4120.

25. Deep, A., Gu, Y., Gao, Y.-Q., Ego, K.M., Herzik, M.A., Zhou, H., and Corbett, K.D. (2022). The SMC-family Wadjet complex protects bacteria from plasmid transformation by recognition and cleavage of closed-circular DNA. Mol. Cell 82, 4145–4159.e7. 10.1016/j.molcel.2022.09.008.

26. Liu, H.W., Roisné-Hamelin, F., Beckert, B., Li, Y., Myasnikov, A., and Gruber, S. (2022). DNA-measuring Wadjet SMC ATPases restrict smaller circular plasmids by DNA cleavage. Mol. Cell 82, 4727–4740.e6. 10.1016/j.molcel.2022.11.015.

27. Bürmann, F., Lee, B.-G., Than, T., Sinn, L., O’Reilly, F.J., Yatskevich, S., Rappsilber, J., Hu, B., Nasmyth, K., and Löwe, J. (2019). A folded conformation of MukBEF and cohesin. Nat. Struct. Mol. Biol. 26, 227–236. 10.1038/s41594-019-0196-z.

28. Lee, B.-G., Merkel, F., Allegretti, M., Hassler, M., Cawood, C., Lecomte, L., O’Reilly, F.J., Sinn, L.R., Gutierrez-Escribano, P., Kschonsak, M., et al. (2020). Cryo-EM structures of holo condensin reveal a subunit flip-flop mechanism. Nat. Struct. Mol. Biol. 27, 743–751. 10.1038/s41594-020-0457-x.

29. Collier, J.E., Lee, B.-G., Roig, M.B., Yatskevich, S., Petela, N.J., Metson, J., Voulgaris, M., Gonzalez Llamazares, A., Löwe, J., and Nasmyth, K.A. (2020). Transport of DNA within cohesin involves clamping on top of engaged heads by Scc2 and entrapment within the ring by Scc3. eLife 9. 10.7554/eLife.59560.

30. Badrinarayanan, A., Reyes-Lamothe, R., Uphoff, S., Leake, M.C., and Sherratt, D.J. (2012). In vivo architecture and action of bacterial structural maintenance of chromosome proteins. Science 338, 528–531. 10.1126/science.1227126.

31. Hopfner, K.P., Karcher, A., Shin, D.S., Craig, L., Arthur, L.M., Carney, J.P., and Tainer, J.A. (2000). Structural biology of Rad50 ATPase: ATP-driven conformational control in DNA double-strand break repair and the ABC-ATPase superfamily. Cell 101, 789–800.

32. Woo, J.-S., Lim, J.-H., Shin, H.-C., Suh, M.-K., Ku, B., Lee, K.-H., Joo, K., Robinson, H., Lee, J., Park, S.-Y., et al. (2009). Structural studies of a bacterial condensin complex reveal ATP-dependent disruption of intersubunit interactions. Cell 136, 85–96. 10.1016/j.cell.2008.10.050.

33. Murayama, Y., and Uhlmann, F. (2015). DNA Entry into and Exit out of the Cohesin Ring by an Interlocking Gate Mechanism. Cell 163, 1628–1640. 10.1016/j.cell.2015.11.030.

34. Karaboja, X., Ren, Z., Brandão, H.B., Paul, P., Rudner, D.Z., and Wang, X. (2021). XerD unloads bacterial SMC complexes at the replication terminus. Mol. Cell 81, 756–766.e8. 10.1016/j.molcel.2020.12.027.

35. Houlard, M., Cutts, E.E., Shamim, M.S., Godwin, J., Weisz, D., Presser Aiden, A., Lieberman Aiden, E., Schermelleh, L., Vannini, A., and Nasmyth, K. (2021). MCPH1 inhibits Condensin II during interphase by regulating its SMC2-Kleisin interface. eLife 10, e73348. 10.7554/eLife.73348.

36. Gruber, S., Arumugam, P., Katou, Y., Kuglitsch, D., Helmhart, W., Shirahige, K., and Nasmyth, K. (2006). Evidence that loading of cohesin onto chromosomes involves opening of its SMC hinge. Cell 127, 523–537. 10.1016/j.cell.2006.08.048.

37. Arumugam, P., Gruber, S., Tanaka, K., Haering, C.H., Mechtler, K., and Nasmyth, K. (2003). ATP hydrolysis is required for cohesin’s association with chromosomes. Curr Biol 13, 1941–1953.

38. Collier, J.E., and Nasmyth, K.A. (2022). DNA passes through cohesin’s hinge as well as its Smc3-kleisin interface. eLife 11, e80310. 10.7554/eLife.80310.

39. Chan, K.-L., Roig, M.B., Hu, B., Beckouët, F., Metson, J., and Nasmyth, K. (2012). Cohesin’s DNA Exit Gate Is Distinct from Its Entrance Gate and Is Regulated by Acetylation. Cell 150, 961–974. 10.1016/j.cell.2012.07.028.

40. Roisné-Hamelin, F., Liu, H.W., Taschner, M., Li, Y., and Gruber, S. (2024). Structural basis for plasmid restriction by SMC JET nuclease. Mol. Cell 84, 883–896.e7. 10.1016/j.molcel.2024.01.009.

41. Vos, S.M., Stewart, N.K., Oakley, M.G., and Berger, J.M. (2013). Structural basis for the MukB-topoisomerase IV interaction and its functional implications in vivo. EMBO J 32, 2950–2962. 10.1038/emboj.2013.218.

42. Wilkinson, M., Wilkinson, O.J., Feyerherm, C., Fletcher, E.E., Wigley, D.B., and Dillingham, M.S. (2022). Structures of RecBCD in complex with phage-encoded inhibitor proteins reveal distinctive strategies for evasion of a bacterial immunity hub. eLife 11, e83409. 10.7554/eLife.83409.

43. Pacumbaba, R., and Center, M.S. (1975). Partial purification and properties of a bacteriophage T7 inhibitor of the host exonuclease V activity. J. Virol. 16, 1200–1207. 10.1128/JVI.16.5.1200-1207.1975.

44. Lin, L. Study of bacteriophage T7 gene 5.9 and gene 5.5.

45. Kimanius, D., Jamali, K., Wilkinson, M.E., Lövestam, S., Velazhahan, V., Nakane, T., and Scheres, S.H.W. (2024). Data-driven regularization lowers the size barrier of cryo-EM structure determination. Nat. Methods 21, 1216–1221. 10.1038/s41592-024-02304-8.

46. Lee, B.-G., Rhodes, J., and Löwe, J. (2022). Clamping of DNA shuts the condensin neck gate. Proc. Natl. Acad. Sci. U. S. A. 119, e2120006119. 10.1073/pnas.2120006119.

47. Muir, K.W., Li, Y., Weis, F., and Panne, D. (2020). The structure of the cohesin ATPase elucidates the mechanism of SMC-kleisin ring opening. Nat. Struct. Mol. Biol. 27, 233– 239. 10.1038/s41594-020-0379-7.

48. Li, S., Yu, Y., Zheng, J., Miller-Browne, V., Ser, Z., Kuang, H., Patel, D.J., and Zhao, X. (2023). Molecular basis for Nse5-6 mediated regulation of Smc5/6 functions. Proc. Natl. Acad. Sci. U. S. A. 120, e2310924120. 10.1073/pnas.2310924120.

49. Li, Q., Zhang, J., Haluska, C., Zhang, X., Wang, L., Liu, G., Wang, Z., Jin, D., Cheng, T., Wang, H., et al. (2024). Cryo-EM structures of Smc5/6 in multiple states reveal its assembly and functional mechanisms. Nat. Struct. Mol. Biol. 10.1038/s41594-024-01319-1.

50. Pradhan, B., Deep, A., König, J., Baaske, M.D., Corbett, K.D., and Kim, E. (2024). Loop extrusion-mediated plasmid DNA cleavage by the bacterial SMC Wadjet complex. BioRxiv Prepr. Serv. Biol., 2024.02.17.580791. 10.1101/2024.02.17.580791.

51. Wang, X., Hughes, A.C., Brandão, H.B., Walker, B., Lierz, C., Cochran, J.C., Oakley, M.G., Kruse, A.C., and Rudner, D.Z. (2018). In Vivo Evidence for ATPase-Dependent DNA Translocation by the Bacillus subtilis SMC Condensin Complex. Mol. Cell 71, 841–847.e5. 10.1016/j.molcel.2018.07.006.

52. Ganji, M., Shaltiel, I.A., Bisht, S., Kim, E., Kalichava, A., Haering, C.H., and Dekker, C. (2018). Real-time imaging of DNA loop extrusion by condensin. Science 360, 102–105. 10.1126/science.aar7831.

53. Ku, B., Lim, J.-H., Shin, H.-C., Shin, S.-Y., and Oh, B.-H. (2010). Crystal structure of the MukB hinge domain with coiled-coil stretches and its functional implications. Proteins 78, 1483– 1490. 10.1002/prot.22664.

54. Dupont, L., Bloor, S., Williamson, J.C., Cuesta, S.M., Shah, R., Teixeira-Silva, A., Naamati, A., Greenwood, E.J.D., Sarafianos, S.G., Matheson, N.J., et al. (2021). The SMC5/6 complex compacts and silences unintegrated HIV-1 DNA and is antagonized by Vpr. Cell Host Microbe 29, 792–805.e6. 10.1016/j.chom.2021.03.001.

55. Isaev, A., Drobiazko, A., Sierro, N., Gordeeva, J., Yosef, I., Qimron, U., Ivanov, N.V., and Severinov, K. (2020). Phage T7 DNA mimic protein Ocr is a potent inhibitor of BREX defence. Nucleic Acids Res. 48, 7601–7602. 10.1093/nar/gkaa510.

56. Kennaway, C.K., Obarska-Kosinska, A., White, J.H., Tuszynska, I., Cooper, L.P., Bujnicki, J.M., Trinick, J., and Dryden, D.T.F. (2009). The structure of M.EcoKI Type I DNA methyltransferase with a DNA mimic antirestriction protein. Nucleic Acids Res. 37, 762– 770. 10.1093/nar/gkn988.

57. Ye, F., Kotta-Loizou, I., Jovanovic, M., Liu, X., Dryden, D.T., Buck, M., and Zhang, X. (2020). Structural basis of transcription inhibition by the DNA mimic protein Ocr of bacteriophage T7. eLife 9, e52125. 10.7554/eLife.52125.

58. Nechaev, S., and Severinov, K. (1999). Inhibition of Escherichia coli RNA polymerase by bacteriophage T7 gene 2 protein. J. Mol. Biol. 289, 815–826. 10.1006/jmbi.1999.2782.

59. Kiro, R., Molshanski-Mor, S., Yosef, I., Milam, S.L., Erickson, H.P., and Qimron, U. (2013). Gene product 0.4 increases bacteriophage T7 competitiveness by inhibiting host cell division. Proc. Natl. Acad. Sci. U. S. A. 110, 19549–19554. 10.1073/pnas.1314096110.

60. Wang, H.-C., Chou, C.-C., Hsu, K.-C., Lee, C.-H., and Wang, A.H.-J. (2019). New paradigm of functional regulation by DNA mimic proteins: Recent updates. IUBMB Life 71, 539–548. 10.1002/iub.1992.

61. Wang, H.-C., Ho, C.-H., Hsu, K.-C., Yang, J.-M., and Wang, A.H.-J. (2014). DNA mimic proteins: functions, structures, and bioinformatic analysis. Biochemistry 53, 2865–2874. 10.1021/bi5002689.

62. Millman, A., Bernheim, A., Stokar-Avihail, A., Fedorenko, T., Voichek, M., Leavitt, A., Oppenheimer-Shaanan, Y., and Sorek, R. (2020). Bacterial Retrons Function In Anti-Phage Defense. Cell 183, 1551–1561.e12. 10.1016/j.cell.2020.09.065.

63. Davidson, I.F., Bauer, B., Goetz, D., Tang, W., Wutz, G., and Peters, J.-M. (2019). DNA loop extrusion by human cohesin. Science 366, 1338–1345. 10.1126/science.aaz3418.

64. Pradhan, B., Barth, R., Kim, E., Davidson, I.F., Bauer, B., van Laar, T., Yang, W., Ryu, J.-K., van der Torre, J., Peters, J.-M., et al. (2022). SMC complexes can traverse physical roadblocks bigger than their ring size. Cell Rep. 41, 111491. 10.1016/j.celrep.2022.111491.

65. Bauer, B.W., Davidson, I.F., Canena, D., Wutz, G., Tang, W., Litos, G., Horn, S., Hinterdorfer, P., and Peters, J.-M. (2021). Cohesin mediates DNA loop extrusion by a “swing and clamp” mechanism. Cell 184, 5448–5464.e22. 10.1016/j.cell.2021.09.016.

66. Srinivasan, M., Scheinost, J.C., Petela, N.J., Gligoris, T.G., Wissler, M., Ogushi, S., Collier, J.E., Voulgaris, M., Kurze, A., Chan, K.-L., et al. (2018). The Cohesin Ring Uses Its Hinge to Organize DNA Using Non-topological as well as Topological Mechanisms. Cell 173, 1508–1519.e18. 10.1016/j.cell.2018.04.015.

67. Tedeschi, A., Wutz, G., Huet, S., Jaritz, M., Wuensche, A., Schirghuber, E., Davidson, I.F., Tang, W., Cisneros, D.A., Bhaskara, V., et al. (2013). Wapl is an essential regulator of chromatin structure and chromosome segregation. Nature. 10.1038/nature12471.

68. Higashi, T.L., Eickhoff, P., Sousa, J.S., Locke, J., Nans, A., Flynn, H.R., Snijders, A.P., Papageorgiou, G., O’Reilly, N., Chen, Z.A., et al. (2020). A Structure-Based Mechanism for DNA Entry into the Cohesin Ring. Mol. Cell 79, 917–933.e9. 10.1016/j.molcel.2020.07.013.

69. Wilkinson, M., Chaban, Y., and Wigley, D.B. (2016). Mechanism for nuclease regulation in RecBCD. eLife 5, e18227. 10.7554/eLife.18227.

70. Zürcher, J.F., Kleefeldt, A.A., Funke, L.F.H., Birnbaum, J., Fredens, J., Grazioli, S., Liu, K.C., Spinck, M., Petris, G., Murat, P., et al. (2023). Continuous synthesis of E. coli genome sections and Mb-scale human DNA assembly. Nature 619, 555–562. 10.1038/s41586-023-06268-1.

71. Engler, C., Kandzia, R., and Marillonnet, S. (2008). A one pot, one step, precision cloning method with high throughput capability. PLoS One 3, e3647. 10.1371/journal.pone.0003647.

72. Butt, T.R., Edavettal, S.C., Hall, J.P., and Mattern, M.R. (2005). SUMO fusion technology for difficult-to-express proteins. Protein Expr Purif 43, 1–9. 10.1016/j.pep.2005.03.016.

73. Studier, F.W. (2005). Protein production by auto-induction in high density shaking cultures. Protein Expr Purif 41, 207–234.

74. Cronan, J.E. (2006). A family of arabinose-inducible Escherichia coli expression vectors having pBR322 copy control. Plasmid 55, 152–157. 10.1016/j.plasmid.2005.07.001.

75. Goedhart, J., and Luijsterburg, M.S. (2020). VolcaNoseR is a web app for creating, exploring, labeling and sharing volcano plots. Sci. Rep. 10, 20560. 10.1038/s41598-020-76603-3.

76. Steinegger, M., and Söding, J. (2017). MMseqs2 enables sensitive protein sequence searching for the analysis of massive data sets. Nat. Biotechnol. 35, 1026–1028. 10.1038/nbt.3988.

77. Katoh, K., and Standley, D.M. (2013). MAFFT multiple sequence alignment software version 7: improvements in performance and usability. Mol. Biol. Evol. 30, 772–780. 10.1093/molbev/mst010.

78. Steinegger, M., Meier, M., Mirdita, M., Vöhringer, H., Haunsberger, S.J., and Söding, J. (2019). HH-suite3 for fast remote homology detection and deep protein annotation. BMC Bioinformatics 20, 473. 10.1186/s12859-019-3019-7.

79. Jumper, J., Evans, R., Pritzel, A., Green, T., Figurnov, M., Ronneberger, O., Tunyasuvunakool, K., Bates, R., Žídek, A., Potapenko, A., et al. (2021). Highly accurate protein structure prediction with AlphaFold. Nature 596, 583–589. 10.1038/s41586-021-03819-2.

80. Bürmann, F., Basfeld, A., Vazquez Nunez, R., Diebold-Durand, M.-L., Wilhelm, L., and Gruber, S. (2017). Tuned SMC Arms Drive Chromosomal Loading of Prokaryotic Condensin. Mol. Cell 65, 861–872.e9. 10.1016/j.molcel.2017.01.026.

81. Minh, B.Q., Schmidt, H.A., Chernomor, O., Schrempf, D., Woodhams, M.D., von Haeseler, A., and Lanfear, R. (2020). IQ-TREE 2: New Models and Efficient Methods for Phylogenetic Inference in the Genomic Era. Mol. Biol. Evol. 37, 1530–1534. 10.1093/molbev/msaa015.

82. Letunic, I., and Bork, P. (2024). Interactive Tree of Life (iTOL) v6: recent updates to the phylogenetic tree display and annotation tool. Nucleic Acids Res. 52, W78–W82. 10.1093/nar/gkae268.

83. Scheres, S.H.W. (2012). RELION: implementation of a Bayesian approach to cryo-EM structure determination. J. Struct. Biol. 180, 519–530. 10.1016/j.jsb.2012.09.006.

84. Rohou, A., and Grigorieff, N. (2015). CTFFIND4: Fast and accurate defocus estimation from electron micrographs. J. Struct. Biol. 192, 216–221. 10.1016/j.jsb.2015.08.008.

85. Wagner, T., Merino, F., Stabrin, M., Moriya, T., Antoni, C., Apelbaum, A., Hagel, P., Sitsel, O., Raisch, T., Prumbaum, D., et al. (2019). SPHIRE-crYOLO is a fast and accurate fully automated particle picker for cryo-EM. Commun. Biol. 2, 218. 10.1038/s42003-019-0437-z.

86. Punjani, A., Rubinstein, J.L., Fleet, D.J., and Brubaker, M.A. (2017). cryoSPARC: algorithms for rapid unsupervised cryo-EM structure determination. Nat. Methods 14, 290–296. 10.1038/nmeth.4169.

87. Pettersen, E.F., Goddard, T.D., Huang, C.C., Meng, E.C., Couch, G.S., Croll, T.I., Morris, J.H., and Ferrin, T.E. (2021). UCSF ChimeraX: Structure visualization for researchers, educators, and developers. Protein Sci. Publ. Protein Soc. 30, 70–82. 10.1002/pro.3943.

88. Rosenthal, P.B., and Henderson, R. (2003). Optimal determination of particle orientation, absolute hand, and contrast loss in single-particle electron cryomicroscopy. J. Mol. Biol. 333, 721–745. 10.1016/j.jmb.2003.07.013.

89. Croll, T.I. (2018). ISOLDE: a physically realistic environment for model building into low-resolution electron-density maps. Acta Crystallogr. Sect. Struct. Biol. 74, 519–530. 10.1107/S2059798318002425.

90. Emsley, P., Lohkamp, B., Scott, W.G., and Cowtan, K. (2010). Features and development of Coot. Acta Crystallogr. D Biol. Crystallogr. 66, 486–501. 10.1107/S0907444910007493.

91. Afonine, P.V., Poon, B.K., Read, R.J., Sobolev, O.V., Terwilliger, T.C., Urzhumtsev, A., and Adams, P.D. (2018). Real-space refinement in PHENIX for cryo-EM and crystallography. Acta Crystallogr. Sect. Struct. Biol. 74, 531–544. 10.1107/S2059798318006551.

92. Cross, S.J., Fisher, J.D.J.R., and Jepson, M.A. ModularImageAnalysis (MIA): Assembly of modularised image and object analysis workflows in ImageJ. J. Microsc. n/a. 10.1111/jmi.13227.

